# SEIZURE OCCURRENCE IN FCD TYPE II IS PREDICTED BY LESION POSITION AND LINKED TO CYTOARCHITECTURAL ALTERATIONS

**DOI:** 10.1101/2025.10.20.683364

**Authors:** N Procházková, CVL Olson, N Shariati, MT Rozlivková, J Populová, M Řehořová, J Kudláček, P Jiruška, O Novák

## Abstract

Focal cortical dysplasia (FCD) is a common malformation of cortical development and a major cause of early-onset, drug-resistant epilepsy. FCD type II is defined by abnormal lamination, altered cellular composition, and pathological cells, notably dysmorphic neurons (DNs) and balloon cells. DNs are thought to drive epileptogenicity through both cell-autonomous and non-cell-autonomous mechanisms, the latter including not only aberrant connectivity but also indirect modulation of excitability in local cell populations.

We performed a multiscale structural and morphological analysis to elucidate the basis of FCD epileptogenicity and the impact of somatic mTOR mutations during brain development. Using a mouse model of FCD type II, we show that lesions in frontal and motor cortical regions are the strongest predictors of spontaneous seizure occurrence. This localization-dependent epileptogenicity offers an experimental explanation for the higher clinical epileptogenicity of frontal FCDs and suggests that posterior lesions may remain silent—an open question in human pathology. In our model, FCD tissue displayed considerable expansion, with cortical thickness up to ∼20% in seizure-bearing animals. This expansion coincided with an overall ∼40% reduction in neuronal density, consistent with tissue hypertrophy. DN density did not differ between seizure and non-seizure animals, challenging the notion that higher DN load directly predicts epileptogenesis.

At the microscopic level, we describe DN axonal pathologies, including giant varicosities. In the cortex, these appeared as vesicle-filled boutons, whereas along callosal axons they were frequent but largely empty. Bouton density was markedly reduced in FCD cortex. Together, these findings leave the net synaptic effect of dysmorphic neurons unresolved, challenging the assumption that axonal hypertrophy translates into increased excitatory drive.

While morphological abnormalities in FCD type II are well documented, their functional consequences remain incompletely understood. Here, we used macro- and microscopic structural features of FCDII to assess seizure susceptibility, providing new insights into epileptogenesis.

## INTRODUCTION

Focal cortical dysplasia (FCD) is a leading cause of early-onset, drug-resistant epilepsy in both children and young adults [1–3]. It represents one of the most common malformations of cortical development [4–6] and manifests in several clinically distinct subtypes [7–9]. Structural hallmarks of FCD type II include disrupted neuronal migration, cortical dyslamination, and the presence of dysmorphic neurons (DNs), which exhibit aberrant size, shape, and extensive dendritic arborization. While the morphological abnormalities in FCD type II have been extensively investigated, the relationship between structure and its pathological functional consequences is not fully understood.

The molecular basis of FCD type II has been linked to hyperactivation of the PI3K/AKT/mTOR signaling pathway. Pathogenic variants in the genes encoding proteins of this pathway, predominantly including the mutations directly affecting the mechanistic target of rapamycin (*mTOR*) gene, lead to dysregulation of key cellular processes including the metabolism, growth, and protein synthesis [9–11]. Persistent mTOR hyperactivation not only results in cellular hypertrophy of dysmorphic neurons but also disrupts normal cortical cytoarchitecture, including changes in the extracellular matrix [12]. These alterations manifest as cortical thickening, dyslamination, white matter heterotopia, and widespread alterations in synaptic architecture, including dendritic spine loss [13] and imbalanced inhibitory signaling, such as abnormally enhanced perisomatic inhibition from parvalbumin (PV)-positive interneurons [14, 15]. Additionally, structural changes have been reported in long-range axonal projections to the contralateral hemisphere, including abnormally enlarged synaptic boutons [16]. While these abnormalities of callosal axonal projections may contribute to the epileptogenicity of FCD type II, alterations in synaptic morphology within the lesion itself have not been systematically investigated.

A separate question related to cortical malformations is the relationship between lesion size and location within the brain, and occurrence of spontaneous seizures. Demonstrating a close relationship between these lesion characteristics and specific cortical functional areas could provide predictive value, for example by helping determine the extent and location of cortical resection in epilepsy surgery [15, 17].

Animal models, especially those employing *in utero* electroporation (IUE) to introduce pathogenic variants of mTOR pathway genes, have proven instrumental in recapitulating key histological and electrophysiological features of human FCD type II [18]. However, while advances in imaging and genetic techniques have deepened our understanding of the role of mTOR pathway hyperactivation in FCD, the relationship between lesion structure and the occurrence of spontaneous seizures is still not fully elucidated. Contributing factors likely include lesion location and size, cellular heterogeneity, differences in neuronal cell counts (including the proportion of mutation-carrying neurons), and alterations in synaptic architecture.

In this study, we expand current knowledge of how both macroscopic and microscopic structural features of FCD type II lesions relate to seizure susceptibility. Using a comprehensive morphological analysis in a mouse IUE model, we characterize lesion properties at both whole-brain and cellular resolution. At the macroscopic level, we map the spatial extent and localization of FCD lesions. At the microscopic level, we quantify DNs, interneuron subtypes, and long-range axonal projections. We then correlate all structural parameters with the incidence of spontaneous seizures.

## MATERIALS AND METHODS

### Sample preparations

#### Animals

Animals of either sex were housed in groups under standard conditions in a room with controlled temperature (22 ± 1 °C) and a 12/12 h light/dark cycle. Experimental mice were generated by crossing SST-IRES-Cre (Cre recombinase expressed under the somatostatin [SST] promoter; Jackson Laboratory, #018973) or PV-P2A-Cre (Cre recombinase expressed under the parvalbumin [PV] promoter; Jackson Laboratory, #012358) mice with Ai14 reporter mice (Cre-dependent expression of the red fluorescent protein tdTomato; Jackson Laboratory, #007909). In some experiments, wild-type (WT) C57BL/6J mice (Jackson Laboratory Cat. No. #000664) were used. The study involved 43 young adult animals of both sexes, all of which underwent the full experimental pipeline outlined in Fig. 1A and described in *Methods*. In SST-IRES-Cre/flex-tdTomato and PV-P2A-Cre/flex-tdTomato genotypes, we prepared mice with FCD (n = 13 and n = 10, respectively) and two types of controls in which the neurons were transfected with a plasmid carrying a gene for either EGFP only (n = 4 and n = 4, respectively), or a wild-type allele of the *mTOR* gene (n = 3 and n = 1, respectively). All C57BL/6J, which underwent the full experimental pipeline (n = 8, Fig. 1A), were electroporated with a plasmid mixture that included a plasmid with mutated *mTOR* as described in Fig. 1 and below in *Materials and Methods*. Additionally, we used three C57BL/6J mice in which the protocol included only IUE of a plasmid mixture and a later injection of an adenosine-associated virus (AAV) into the FCD lesion at week 12. These mice were not implanted with electrodes and were transcardially perfused at week 15 and used only for mounted brain slice preparation (see below). In these mice the plasmid mixture contained a plasmid with mutated *mTOR* followed by a Cre recombinase and a different lesion-labeling plasmid, CAG-iRFP_713_.

**Fig. 1.**
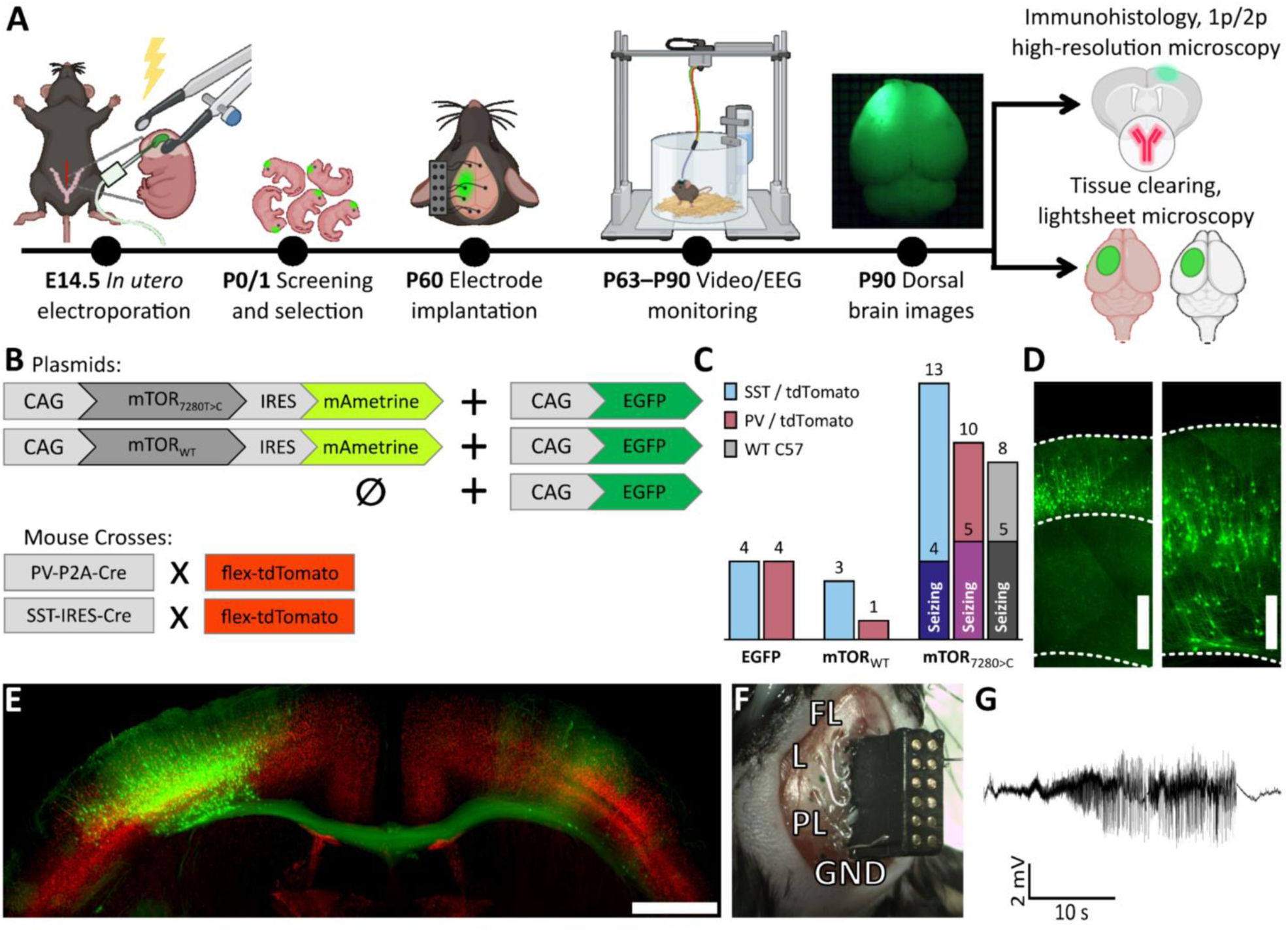
Preparation and characterization of the FCD type II mouse model. (**A**)Schematic overview of the experimental pipeline. (**B**) Plasmid constructs used to generate experimental and control groups, along with breeding schemes for producing the final mouse genotypes. Experimental animals were electroporated with a construct carrying the mutated *mTOR* gene co-expressing mAmetrine, together with an EGFP-encoding plasmid. The first control group was electroporated with wild-type *mTOR* and EGFP, both co-expressing mAmetrine. The second control group received EGFP alone. Mice were bred on a C57BL/6J background and were either SST-IRES-Cre or PV-P2A-Cre crossed with flex-tdTomato, or wild-type C57BL/6J animals. (**C**) Final numbers of mice used in the complete experimental workflow showed in (**A**), subdivided by interneuron subtype (tdTomato labeling) and seizure phenotype. (**D**) Representative images of GFP+ cell distribution across cortical depth in control and FCD animals; scale bar = 500 μm. (**E**) Representative coronal brain section from a mutant FCD PV-tdTomato mouse, showing long-range axonal projections from the FCD lesion to the contralateral homotopic area, where green represents GFP and red represents tdTomato fluorescence; scale bar = 1 mm. (**F**) Image of an implanted EEG connector and electrodes. FL, frontal left; L, lesion; PL, perilesional; GND, ground. Additional electrodes (frontal right [FR], contralateral homotopic [C], and reference) were also implanted but are visible in the image. (**G**) Example EEG trace showing a seizure recorded from the lesion (L) electrode.

All experiments were performed under the Animal Care and Animal Protection Law of the Czech Republic, fully compatible with the guidelines of the European Union directive 2010/63/EU. The protocol was approved by the Ethics Committees of the Second Faculty of Medicine (Project License No. MSMT-31765/2019–4).

#### *In utero* electroporation and FCD presence evaluation

The FCD lesion was induced by IUE [11, 19, 20]. Pregnant mice (day 14.5 ± 0.5 post-fertilization) were anesthetized with isoflurane (5 % induction, 1.5–2 % maintenance), subcutaneously injected with ketoprofen (5 mg/kg), and positioned on their backs. A midline incision was performed along the *linea alba*. The uterine horns were gently exposed. Using pulled and sharply beveled glass capillaries, the left lateral ventricle of each embryo was injected with a mixture of plasmids dissolved in ACSF containing 2.0 μg/ml Fast Green (F7252, Sigma, USA). In FCD model animals, the plasmid mixture contained 3.0 μg/μl mutant *mTOR* (c.7280T>C, SoVarGen Co., Ltd., Korea) under the control of the CAG promoter; these mice are further termed as FCD animals. The m*TOR* gene was followed by a gene encoding a fluorescent protein mAmetrine, however, we were not able to detect mAmetrine fluorescence and we thus do not consider this fluorescent protein in this work. For the lesion visualization, the *mTOR* plasmid was mixed with an expression plasmid CAG-EGFP at a final concentration of 1.5 μg/μl.

This work contains two types of control animals. One group of animals was electroporated with a wild-type *mTOR* gene (3.0 μg/μl) in the mixture with 1.5 μg/μl CAG-EGFP. In the second group of control animals, we electroporated 1.5 μg/μl CAG-EGFP only. After each injection, the electroporation was performed by positioning forceps-type electrodes (3 mm platinum plated pads, CUY650P3, Nepagene) on the head of the embryo, and applying five 35–45V, 50 ms pulses with 950 ms interpulse intervals using a custom–made high-resolution electroporator with a 40 mA current limiter [20]. After electroporation of the pups, the uterine horns were returned to the abdominal cavity and the wound was sutured in individual layers. The mouse was turned on its abdomen and kept on the inhalation mask with pure oxygen until regaining consciousness. Then, the mouse was returned to its home cage that was placed on a heating pad overnight. Following the surgery, the mouse was daily injected with ketoprofen (5mg/kg) for two consecutive days. When the pups were born, we screened them for fluorescence using a strong, collimated blue LED (GT-P04B3410320, GETIAN, China) with a band-pass excitation filter (475/50 nm, Edmund Optics, USA) and a stereomicroscope (Model STM 823 N, Optika, Italy) with a large-diameter lens cap containing a green band-pass filter (525/50 nm, Edmund Optics, USA). Only the successfully electroporated pups in which we detected a properly positioned green fluorescent spot were kept in the experiment. Animals with heterogeneous or bilateral fluorescent lesion patterns were excluded.

#### Electrode implantation surgery

The surgery was performed in animals at 8–10 weeks of age. Mice were anesthetized with isoflurane (5 % induction, 1.5–2 % maintenance), the fur on top of the animal’s head was shaved, and a small area of skin was removed to expose the dorsal skull. We gently removed the soft tissues overlying the bone. With the skull kept wet and relatively transparent, we used the strong blue LED and the green bandpass filter cap (525/50 nm, #86-963, Edmund Optics, USA) on the objective of the surgical microscope to map the fluorescently labeled lesion onto the skull hallmarks (bregma, lambda). Telecentric fluorescent images of the dorsal skull were further used for morphometry in this project. Animals in which we detected a heterogeneous or bilateral lesion in this adult stage were excluded from the experiment. The position of the lesion was used to properly place the recording electrodes. Custom EEG implants were prepared by soldering insulated silver wires (127 μm in diameter, AM Systems, Inc., USA, Cat. no. 786000) to prefabricated connectors (TME Electronic Components, Poland, Cat. no. DS1065-03-2*6S8BV). Seven holes, adjusted to the position of the GFP signal, were drilled through the skull using a high-speed microdrill (Success40, Osada, Japan). Two electrodes were placed over the center of the lesion and on its margin, and one electrode was implanted to the contralateral position that was homotopic to the center of the lesion. Two electrodes were implanted over the left and right frontal cortices. The grounding/reference electrodes were placed over the inferior colliculus/cerebellum. All electrodes had coil-shaped endings that were placed on the dura to prevent them from inflicting tissue damage to the brain. The holes were sealed with bone wax (SMI, Belgium, Cat. no. Z046). All the leads together with the connector were attached to the skull using a gel form of cyanoacrylate glue (Loctite, USA, Cat. no. 1363589) cured with a glue accelerator (Insta-Set; Bob Smith Industries, Atascadero, CA, USA). The skin margins were secured and sealed to the connector construct and the skull also using a cyanoacrylate glue.

#### AAV injection surgery

Mice were anesthetized with isoflurane (5 % induction, 1.5–2 % maintenance), the fur on top of the animal head was shaved, and a small cut in the skin over the left parietal cortex was made. We used a strong red laser pointer (50mW, 632 nm), an infrared bandpass filter cap (725/50 nm, #86-967, Edmund Optics, USA) on the objective of the surgical microscope, and an attached NIR-sensitive camera (acA1300-60gmNIR, Basler AG, Germany) to evaluate the position of the lesion (these mice were co-electroporated with CAG-iRFP_713_ instead of CAG-EGFP). We made a 0.5mm hole through the skull using a high-speed microdrill (Success40, Osada, Japan). Using a pulled and beveled (outer diameter 10 µm) glass capillary and a hydraulic injector (capillary: 3-000-203-G-X, injector: NanojectIII, Drummond Scientific, USA), we injected 300 nl of AAV (phSyn1(S)-FLEX-tdTomato-T2A-SypEGFP-WPRE, 51509-AAV1, Addgene, USA) into the cortex; depth −300 µm, titer was 2×10^12^ GC/ml, injection speed 1 nl/s. After the retraction of the capillary, we sealed the hole with a cyanoacrylate glue and sutured the cut in the skin. The mice were perfused for immunohistochemistry (IHC) after three weeks.

#### Immunohistochemistry

After the completion of EEG monitoring, mice were overdosed with intramuscular injection of a mixture of ketamine/xylazine 120 mg/kg and 20 mg/kg, respectively. The animals were transcardially perfused with cold saline and 4% paraformaldehyde (PFA). Brains were collected, post-fixed in PFA overnight, and then stored in PBS with 0.05% azide. To further map the position and quantify the spatial extent of the lesion, the post-fixed brains were photographed using the aforementioned blue excitation/green emission add-on to the surgery microscope. The fixed brains were cut on a vibratome (Leica VT1200 S, Leica, USA) into 60 μm thick coronal slices. The free-floating slices were washed with phosphate saline buffer (PBS) and non-specific binding sites were blocked using ChemiBLOCKER (Sigma-Aldrich, USA). IHC was performed using primary antibodies for neuronal nuclear protein (NeuN; 1:1000, rabbit, Sigma-Aldrich, Cat. no. ABN78, USA), and phospho-S6 ribosomal protein (Ser235/236) (1:600, rabbit, Cell Signalling, Cat. no. #2211, USA). In several slices from the WT C57BL/6J mice, we applied primary antibodies against PV (1:100, rabbit, Abcam, UK, Cat. no. ab181086), synapsin-1 (1:100, rabbit, Sigma-Aldrich, USA, Cat. no. SAB4502903), vesicular glutamate transporter 1 (VGLUT1; 1:100, rabbit, Sigma-Aldrich, USA, Cat. no. AB5062P), and postsynaptic density protein 95 (PSD95; 1:200, goat, Abcam, UK, Cat. No. ab12093). All primary antibodies were incubated with the slices overnight at 4 °C. For the first three primary antibodies, which were raised in rabbit, we used the secondary antibody conjugated with a fluorescent dye CF™ 647 (1:500, anti-rabbit, Sigma-Aldrich, USA, Cat. no. SAB4600185). For the goat PSD95 primary antibody, we used the secondary antibody conjugated with a fluorescent dye Alexa Fluor™ 594 (1:500, anti-goat, Abcam, UK, Cat. No. ab150132). Secondary antibodies were also incubated overnight at 4 °C. After washing, slices were mounted using Fluoromount-G™ Mounting Medium (Fisher Scientific International, Inc., Cat. no. 00-4959-52, USA). We prepared at least two slices from each mouse and each antibody.

#### Brain tissue clearing

Brains that were subject to light-sheet microscopy were treated by the clearing procedure using the CUBIC protocol [21]. We used the protocol for simple immersion and followed the whole process as described in Susaki et al [21]. Briefly, the brains were dissected and post-fixed in 4% PFA for one day at 4 °C. Afterward, they were washed in PBS for at least one day, with the solution changed every 12 hours. The brain samples were delipidated using a CUBIC-1 solution for 1–2 weeks at room temperature, with the solution changed every third day. Following this step, the samples were washed in PBS for one day, with the solution changed every three hours. Finally, the brains were transferred to the refractive index-matching solution CUBIC-2 for five days, with the solution replaced daily.

### Data acquisition

#### EEG recording

Following a five-day recovery period after the surgery, the animals were individually video-EEG monitored continuously for four weeks. A custom-made video-EEG unit consisted of a counterbalance mechanism, slip rings with an active motorized commutator and a video camera. The connector on the head of the animal was connected to a headstage in which the spontaneous electrographic activity was amplified, band-pass filtered (0.1 Hz–1.6 kHz), and sampled at 5 kHz using a 32-channel headstage amplifier with an AD converter (Intan Technologies, USA, chip Cat. no. RHD2132). The data were recorded using an Interface Board (Intan Technologies, USA, Cat. no. C3100) controlled by Spike2 software (10.02, Cambridge Electronic Design, Cambridge, UK). Manual seizure labeling and EEG data analysis were performed using custom-made scripts in a MATLAB 2019b computing environment (The MathWorks, Inc., Natick, Massachusetts, USA).

To analyze interictal activity, we detected spikes (putative interictal epileptiform discharges) from the electrode positioned within the FCD lesion. Spike detection was performed using an automated algorithm based on thresholding of the analytic signal amplitude [22]. The detector was applied with its original parameters and, additionally, with the low-pass filter cutoff frequency reduced to 40 Hz and 30 Hz to verify that the results were independent of specific detector settings. For each mouse, we calculated the average spike rate and compared it across three groups: non-FCD controls, FCD mice without seizures, and FCD mice with seizures.

#### Imaging of lesion cytoarchitecture and projections

Cellular architecture and morphology of GFP-expressing cells, and IHC-labeled NeuN-, pS6-, or PV-positive cells were imaged and recorded using a high-resolution custom-built two-photon microscope equipped with a water-immersion 20×/1.0 NA XLUMPLFLN objective (Olympus, Japan). To excite both EGFP and the CF^™^ 647 fluorescent dye, we delivered femtosecond pulses of 860 nm wavelength using a Chameleon Discovery tunable femtosecond laser (Coherent, USA).

To control the microscope, we used the ScanImage software bundle (MBF Bioscience, USA) and added custom-made scripts that enabled setting the microscope to automatically perform sequential Z-stack imaging at predefined positions. We used a water-immersion 20×/1.0 NA XLUMPLFLN objective (Olympus, Japan) with a 2.0 mm working distance. The 2D field of view (FOV) of each Z-stack was 810×810 μm and was covered by 1024×1024 pixels, with 1 μm Z-step. In each slice, we programmed the microscope to perform many sequential Z-stacks (tiles) to continuously cover the entire area of the lesion and its contralateral projection areas. Individual tiles were separated by 700 μm providing 110 μm overlaps for later stitching.

#### Confocal Microscopy

To image very fine ipsilateral and contralateral axonal projections with synaptic boutons of the transfected neurons, images from coronal brain slices were captured using either a high-resolution scanning confocal microscope, Leica Stellaris 5 (Leica Microsystems, Germany), or a spinning-disk confocal, Andor Dragonfly 503 (Oxford Instruments, UK). The Stellaris 5 microscope was equipped with a HC PL APO 63×/1.40 NA oil objective, HyD-S hybrid detector, and white light laser with pulse picker set to 490 nm (Leica Microsystems, Germany). The final confocal voxel tiles from the Stellaris 5 were sampled at 0.172×0.172×0.202 μm voxels. The Dragonfly 503 microscope was equipped with a HCX PL APO 63×/1.40-0.6 NA oil objective (Leica Microsystems, Germany), a quad excitation dichroic blocking filter set (405 nm/488 nm/561 nm/640 nm), and a Zyla 4.2 Plus sCMOS camera (Oxford Instruments, UK). We imaged in the green channel (Ex: 488 nm laser, Em: BP 525/40 nm). The final confocal voxel tiles from the Dragonfly 503 were sampled at 0.108×0.108×0.220 μm voxels.

In each slice, we performed imaging of six Z-stacks. Three of the Z-stacks were located in the middle of the lesion and were performed at the cortical depth with the upper edge of the FOV corresponding with 1/3, 2/3, and 3/3 of the cortical gray matter thickness in the middle of the lesion. We further termed these locations as L2/3 FOV, L5 FOV, and WM FOV, respectively. Three more Z-stacks were performed in the contralateral area covered with the projections of the FCD neurons. Furthermore, we scanned the coronal brain slices stained for synapsin-1, VGLUT1, and PSD95 to capture the axonal varicosities and confirm their identity. We acquired representative Z-stacks spanning the thickness of the slice, with ROIs located in the corpus callosum and contralateral homotopic area. We performed the scanning in high resolution using the aforementioned HCX PL APO 63×/1.40-0.6 NA oil objective (Leica Microsystems, Germany). We oversampled the theoretical resolution of the microscope with sampling steps at *XY* pixel size <0.1 um and Z-step 0.20 μm for subsequent spatial deconvolution (Richardson-Lucy, Gaussian PSF σ_x,y_ = 300 nm, σ_z_ = 900 nm, 10 iterations).

#### Light-sheet microscopy

Besides obtaining data from IHC using the two-photon microscope, a randomly selected portion of the brains was chosen to undergo light-sheet microscopy to get the full spatial context of FCD lesions. We used a LightSheet Z.1 microscope equipped with a 5×/0.1 NA illumination objective (Zeiss, Germany), a quad excitation dichroic blocking filter set (405 nm/488 nm/561 nm/640 nm), and a PCO.Edge 5.5 sCMOS camera (Excelitas, USA) to scan the entire lesion together with its complete projections. We imaged in green (Ex: 488 nm laser, Em: BP 525/20 nm) and red (Ex: 561 nm laser, Em: BP 595/20 nm) channels to record the morphology and projections of the transfected neurons (GFP) and the numbers and morphology of interneurons (tdTomato). A 5×/0.16 NA objective (Zeiss, Germany) was used for the detection and volumetric imaging. The individual tiles were deconvolved and stitched in Huygens Professional software (24.04, Scientific Volume Imaging, NL, http://svi.nl). The final 3D brain volumes were sampled at 1.17×1.17×4.94 μm voxels.

### Image data processing

#### Macroscopic images of the dorsal skull and the bare explanted brain

Macroscopic fluorescence images of the dorsal cortex (Fig. 1A, Fig. 2C) were taken either with or without the skull using a camera attached to a surgical microscope with a fluorescence add-on (described in FCD presence evaluation) equipped with a Plan-Apochromat objective. With a working distance of >150 mm, this setup ensured sufficient telecentricity. For each brain, the fluorescently-labeled area was manually delineated and a separate black-white image with the delineated mask was created. The person who delineated the lesions was blind both to the character of the site of transfected neurons (FCD or control) and to any possible epilepsy phenotype associated with the particular brain. As a reference brain map, we used the mouse dorsal-cortex quantitative map (Fig. 2A) published by Matt Kirkcaldie (adopted with permission) [23]. The reference map was simplified and manually warped using a similarity transformation to fit the dorsal image of the brain. We then applied the inverse transformation to the respective image containing the black-white mask. This procedure enabled precise projecting of the areas of the transfected cortex onto the original reference map and their comparison.

**Fig. 2.**
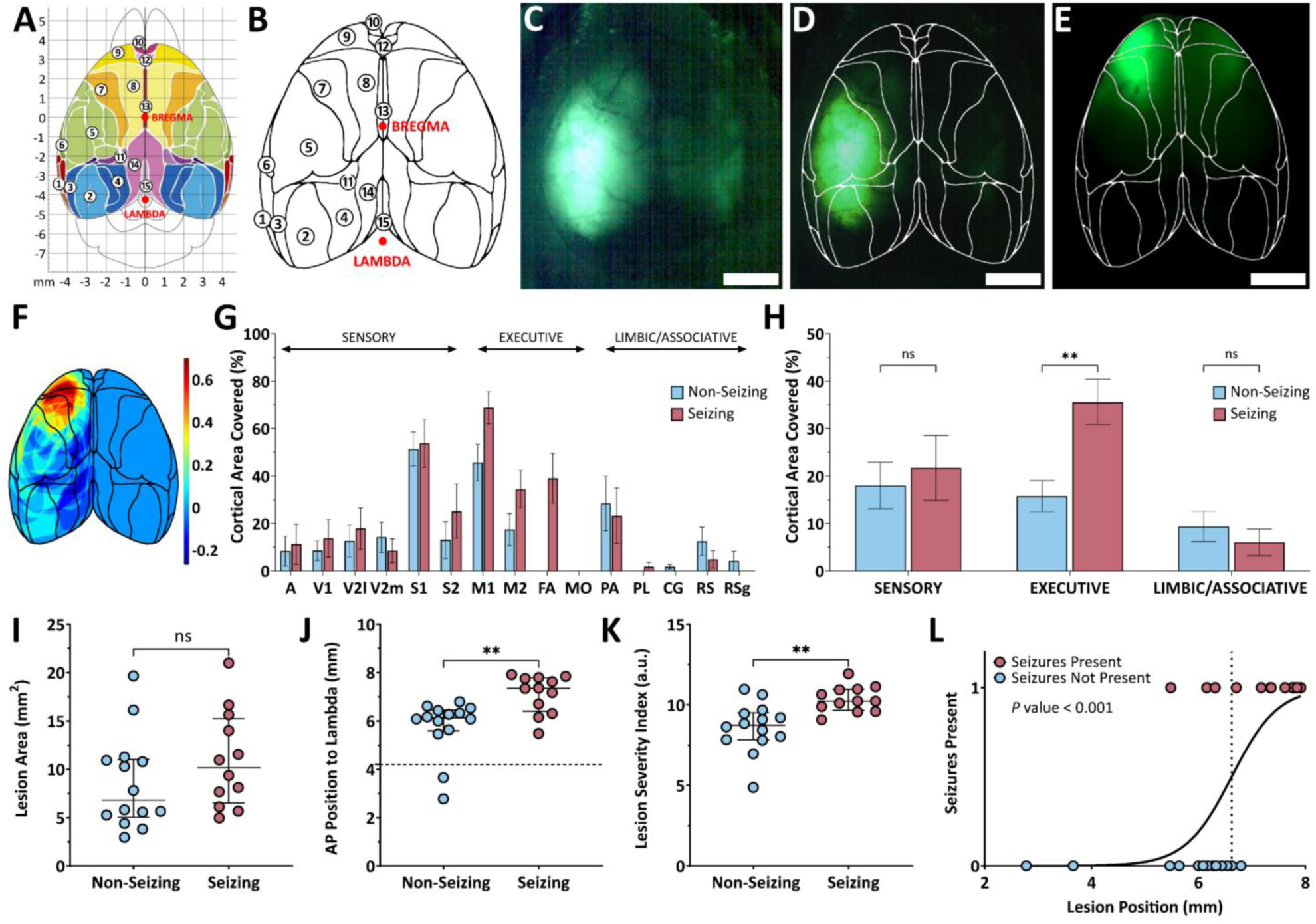
Lesion size and position correlate with seizure occurrence and enable their predictive modeling. **(A)** Schematic map of functional cortical areas in the mouse brain, adapted with permission from [23]. (**B**) The same map simplified to outline the functional areas. (**C**) Dorsal image with a brain with fluorescently-labeled lesion; scale bar = 2 mm. (**D, E**) Examples of brains with fluorescent FCD lesions and overlaid masks of cortical areas; scale bars = 2 mm. (**F**) Heatmap showing spatial overlap of lesions in seizing vs. non-seizing animals. (**G**) Frequency of involvement of cortical areas in FCD lesions of seizing vs. non-seizing mice. (**H**) Summary comparison of lesion involvement in Sensory, Executive, and Limbic/Associative regions; two-way ANOVA with ‘Seizing’ and ‘Brain Area’ as factors (‘Brain Area’ *p* < 0.001). Šidák’s multiple comparisons test: Sensory (*p* = 0.56), Executive (*p* = 0.002), Limbic (*p* = 0.60). Bars represent group means; whiskers indicate SEM. (**I**) Comparison of lesion sizes in seizing vs. non-seizing animals. (**J**) Anterior– posterior (AP) position of the lesion relative to lambda; dashed line indicates bregma position. (**K**) Lesion severity index (LSI), integrating lesion size and position. (**I–K**) Points represent individual animals, central lines = medians, whiskers = IQRs. (**L**) Simple logistic regression between the lesion position and a binary ‘seizing’ = 1 / ‘non-seizing’ = 0 value; dotted line indicates the 50% curve position (6.62 mm). Specific legends and values to Fig. 2G, H are provided in **Table 1**.

**Table 1.** Legend and specific values to Fig. 2.

| Symbol | Brain Area | Abbr. | Mean (SEM) (%) |  | Functional Area | Mean (SEM) (%) |  |
| --- | --- | --- | --- | --- | --- | --- | --- |
|  |  |  | Non-seizing (n=12) | Seizing (n=7) |  | Non-seizing (n=12) | Seizing (n=7) |
| ① | Auditory cortex | **A** | 8.3 (6.2) | 11.3 (8.5) | **Sensory** | 18.0 (4.9) | 21.7 (6.8) |
| ② | Primary visual cortex | **V1** | 8.5 (4.1) | 13.7 (7.9) |  |  |  |
| ③ | Secondary visual cortex - lateral | **V2l** | 12.6 (6.7) | 17.8 (8.8) |  |  |  |
| ④ | Secondary visual cortex - medial | **V2m** | 14.1 (6.4) | 8.5 (5.0) |  |  |  |
| ⑤ | Primary somatosensory cortex | **S1** | 51.4 (7.2) | 53.8 (10.1) |  |  |  |
| ⑥ | Secondary somatosensory cortex | **S2** | 13.0 (7.7) | 25.2 (11.4) |  |  |  |
| ⑦ | Primary motor cortex | **M1** | 45.7 (7.7) | 68.8 (6.8) | **Executive** | 15.8 (3.3) | 35.6 (4.8) |
| ⑧ | Secondary motor cortex | **M2** | 17.4 (6.9) | 34.5 (7.8) |  |  |  |
| ⑨ | Frontal association cortex | **FA** | 0.00 | 39.1 (10.5) |  |  |  |
| ⑩ | Medial orbital cortex | **MO** | 0.00 | 0.00 |  |  |  |
| ⑪ | Parietal association cortex | **PA** | 28.4 (11.6) | 23.3 (11.7) | **Limbic / Associative** | 9.3 (3.2) | 6.0 (2.8) |
| ⑫ | Prelimbic cortex | **PL** | 0.00 | 1.8 (1.8) |  |  |  |
| ⑬ | Cingulate cortex | **CG** | 1.8 (1.0) | 0.00 |  |  |  |
| ⑭ | Retrosplenial cortex | **RS** | 12.4 (6.0) | 4.9 (3.6) |  |  |  |
| ⑮ | Retrosplenial granular cortex | **RSg** | 4.1 (4.1) | 0.00 |  |  |  |

To display the overall difference in average positions of lesions associated and non-associated with seizures, we summed all nonzero elements of the respective images containing registered masks. In seizing brains, the pixels of the mask were assigned a value of +1/12, and all nonzero elements in the respective non-seizing brains were assigned -1/14 (the denominators correspond to the number of brains). For the evaluation of the relationship between the sizes of the lesions and the seizing phenotype, we simply calculated the delineated areas. To evaluate the influence of the lesion position, we compared the positions of the rostral edges of the lesions. The position is referenced to the Lambda position. To evaluate the combined effect of the size and the position of the lesion, we created a variable *Lesion Severity Index*, which is defined as the square root of the lesion area plus the position of the lesion’s frontal edge referenced to the Lambda point. Finally, we used logistic regression as a test for seizure prediction capability of the lesion position; p(x)=1/(1+exp(−(β_0_+β_1_⋅*x*))).

#### Slice-wide images of GFP-, pS6-, PV- and NeuN-labeled neurons recorded via two-photon microscopy

Z-stacks were first deinterleaved into red and green channels and then each channel stack was split into individual 2D frames using FIJI (ImageJ 2.14.0/1.54f) [24]. The cells in each frame were segmented as 2D masks to mark GFP-positive and CF^™^ 647-positive cells. The segmentation was performed with Cellpose 2.0 using a modified method of human-in-the-loop training [25]. First, cell positions in a single frame and channel were approximated with the default *cyto2* model. These masks were then manually corrected, either by adding additional masks to non-segmented cells, deleting masks from regions where no cell signal could be detected, and/or adjusting the size and position of masks for better cell fitting. After repeating this baseline mask and correction process for 10 frames, we used Cellpose’s neural network to train a custom model (*learning_rate*: 0.2, *weight_decay*: 1e-05, *n_epochs*: 50,000). The masking and correction steps were then repeated using the custom model, and training was performed again. In total, this feedback training process was performed 4–5 times for each channel signal to yield our finalized models (GFP, pS6, PV, and NeuN). All of our sampled frames were then batch segmented automatically with Cellpose running in a Jupyter notebook environment (Python 3.11.5, Jupyter 7.0.8) and saved as *.npy* arrays and *.png* masks.

For the evaluation of size and brightness of individual cells, we proceeded as follows: in each frame (1024×1024 pixels, corresponding to 0.79 μm/pixel) of a Z-stack, we found centroids of the segmented cells, divided their coordinates by a factor of 4, rounded them, and registered the centroids as ones in the respective coordinates in 4x smaller matrix (corresponding to 3.2 μm/px). This step was applied to compensate for the possibility of slightly wandering centroids of the same cell captured in different slices. We summed all such matrices from all slices of the Z-stack. We then thresholded the summed matrix (threshold = 4) and counted the non-zero objects in the 2D image (8-connected neighborhood). The threshold was found empirically to find all cells inside the Z-stack except the upper and lower 4–6 μm-thick margins of the Z-stack. This condition was set in order to prevent the neurons on the margins of the potential consecutive Z-stacks from being counted twice. We further took the number of calculated non-zero objects in 2D as the number of centroid clusters (number of neurons to find). Centroids of the cells in the original .*png* masks from Cellpose were summed to a new 1024×1024 matrix (except the centroids from the upper and lower five slices, corresponding to the upper and lower 5 μm margins of the Z-stack). Finally, we applied *k-means* clustering on the non-zero points of this matrix (the number *k* was taken as the previously evaluated number of neurons to be found). This step refined the positions of the neurons. In each cluster (corresponding to one neuron), we found the slice in which the neuron print had the largest area (in approximately spherical objects such a slice corresponds to their middle sections). The respective delineated area of the neuron was fitted with an ellipse and the length of its major axis, its minor axis, and together with the major axis orientation were recorded. In the two channels, we calculated the maximal intensity projections (MIPs) of our Z-stacks and all tiles belonging to a single brain slice were stitched together. The corresponding *XY* positions of individual neurons in separate tiles were translated in *XY* relative to the tiles’ stitched position to yield a total cell mask across each given slice.

From the stitched MIPs, the lesion area, lesion depth, and cortical thickness were determined manually. First, we evaluated the tangential extent of the lesion determined by lines perpendicular to the cortex surface that separated 1 % of the GFP-positive cells on each side. Then, the delineation was closed with the curve going along the surface and the curve going at the boundary between the gray and the white matter. The thickness of the cortex was measured as a perpendicular distance from the cortex surface to the gray/white matter boundary in the center of the GFP-labeled area. We used two methods to determine the signal intensity threshold. For evaluation of the PI3K/AKT/mTOR pathway hyperactivation via strong pS6 immunolabeling, we developed an algorithm that automatically sets the proper threshold based on the assumption that there are two populations of neurons differing in the intensity of labeling—those with hyperactivated PI3K/AKT/mTOR pathway and those with its normal activity that also produce a baseline level of pS6. For all slices containing FCD, we took the stitched MIP image, applied an automatic contrast adjustment stretching the look-up table between the 1^st^ and the 99^th^ percentile, and calculated the mean intensity of individual cells using their centers and fitted ellipse. We constructed a histogram of 256 equally separated bins (0–255) and calculated a bin count (Suppl. Fig. 2). We fitted the bin count with a bi-Gaussian distribution. The final threshold for the mean brightness of highly ps6-positive cells was set as the mean value of the centers of the two Gauss distributions. We used this method to find the threshold in all animals with FCD. The second method to evaluate the ps6-positive cells was based on the level of nonspecific labeling and autofluorescence of the slice (taken as background). The representative background fluorescence value was counted as a mean intensity over 20 manually selected points in the first cortical layer. Then the threshold was taken as 3× background intensity value. The thresholds obtained using the second method differed from the respective threshold values obtained using the bi-Gaussian method only within a small difference range (±5 %). We developed the second method for use in slices from control animals, where the bi-Gaussian method cannot, by definition, be applied. In GFP and NeuN labeling, we applied the first-layer intensity-related threshold to remove the cells that could be detected only based on their nonspecific autofluorescence. In this case, we set the threshold to value 1.5× first-layer intensity. In case of GFP-positive neurons, we selected the points in the first cortical layer outside the lesion (in GFP excluding the labeled apical dendrites).

#### Axonal boutons analysis

The confocal Z-stacks were analyzed using Imaris 10.2 (Oxford Instruments, UK). Within each stack, nine axons were randomly selected from those axons crossing the middle slices of the Z-stacks and manually traced in 3D using the *Filaments* function. Once the traces were properly centered and their diameters calculated, we exported the trace lengths and diameters for bouton analysis in Matlab. The thicknesses of the axonal shafts were taken as the 5^th^ percentile of the diameter along the length of the axon portion. Relative changes in axon diameter (Suppl. Fig. 4) were analyzed using the *findpeaks* function with the following parameters (fraction of diameter value): minimum peak height = 0.2, minimum peak prominence = 0.1, minimum distance of two accepted peaks = 1.0, peak width = ‘halfheight’. Bouton volumes were calculated as volumes of rotational ellipsoid with *a* = *b* taken as diameters and *c* given by peak width.

### Light-sheet microscopy data

#### Depth profile analysis

In order to assess the depth profile and density from the light-sheet images, we first had to define the surface of each brain. To do this, we calculated the position of all tdTomato cells in our image using the less computationally demanding *Spots* function, which does not return volumetric data, but does return a set of *X*, *Y*, and *Z* coordinates for each detection. From these positions, we created a convex hull around each brain and computed a perpendicular axis in the center of the lesion (taken as center of mass based on the position of individual GFP+ cells). Along this axis, we constructed a 1-mm-wide square column of and calculated the number of cells whose centers belonged to a space given by this column and a 0.01 mm step. The depth distributions were calculated and normalized across depth and to unit sum.

### Statistical analyses

Statistical tests were performed in either MATLAB (R2024b, The MathWorks, Inc., Natick, Massachusetts, USA), Python (3.11.5-Van Rossum, G., & Drake, F. L. (2009), Python 3 Reference Manual. Scotts Valley, CA: CreateSpace), or GraphPad Prism (10.4.1 for Windows, GraphPad Software, Boston, Massachusetts, USA, www.graphpad.com). Values are reported as median (IQR) unless otherwise noted. Nonparametric testing was used when data did not display normality of residuals or non-equal variance due to the low sample numbers. Bonferonni corrections for multiple comparisons were always applied. *P*-values were taken as significant if less than 0.05 (*), 0.01 (**), and 0.001 (***).

## RESULTS

### Overview of the experimental pipeline and animal cohorts

In this experiment, we followed the pipeline illustrated in Figure 1A and thoroughly described in *Materials and Methods*. The design of experimental and control groups is summarized in Fig. 1B. In total, the main part of the study including IUE involved 8 GFP-positive (GFP+) control animals, 4 WT-mTOR animals, and 31 FCD animals. The experimental groups are further stratified based on the tdTomato-labeled interneuron subtype (Fig. 1C). At 8 weeks of age, animals underwent surgery. At the beginning of the surgery, a macroscopic fluorescence image of the brain was acquired through the wet skull. Subsequently, the animals were implanted with a connector and five EEG recording electrodes (Fig. 1F). Each animal was continuously monitored using video-EEG for a minimum of four weeks. Afterward, the animals were sacrificed and fluorescence images of the explanted brains were obtained. Brains were subsequently processed for either immunohistochemistry or tissue clearing, as the two methods are mutually exclusive. As previously reported in this model [11, 26], the brains displayed typically altered cytoarchitecture of the FCD type II (Fig. 1D, E), including the presence of cytomegalic DNs, dyslamination, and white matter heterotopia. EEG recordings revealed spontaneous seizures in 45% of the animals (Fig. 1C, G). In the group of FCD animals with spontaneous seizures, we report the median frequency of 2.6 (1.2–4.4) seizures per day and the median seizure duration 49.3 (40.1–67.1) s (Suppl. Fig. 1 A, B). We analyzed signals from electrodes placed above the type II FCD lesion or the corresponding cortical area in controls (Suppl. Fig. 1C). FCD mice with seizures exhibited a markedly higher rate of interictal epileptiform discharges 21.9 (15.5-35.7) IEDs/min compared with both controls 3.2 (2.2-5.3) IEDs/min and non-seizing FCD mice 5.1 (3.3-6.1) IEDs/min, as assessed by the Mann–Whitney U test (p < 0.001 and p = 0.0016, respectively). The moderate increase observed in non-seizing FCD mice relative to controls was not statistically significant (p = 0.29, Bonferroni-corrected, n = 3).

### Rostro-caudal position and the coverage of the frontal cortical areas represent the main predictive factor of seizure occurrence

Studies utilizing the IUE-based FCD II model have shown that only a subset of FCD mice develop spontaneous seizures [27, 28], suggesting that seizure susceptibility is influenced by additional factors. These may involve the proportion of neurons carrying the mutation within the lesion, the spatial extent of the lesion and its location. We first focused on the latter two parameters. In addition, we stratified each lesion according to the functionally defined cortical areas (Fig. 2A), using a simplified version of a published brain map from [23], in which adjacent secondary areas were merged and the major cranial landmarks—bregma and lambda—were highlighted (Fig. 2B).

To quantify lesion size and position, we overlaid the simplified map onto lesion fluorescence macroscopic dorsal images taken either at the beginning of surgery (Fig. 2C) or after brain explantation at the end of the *in vivo* protocol. We used either the bregma/lambda coordinates or overall brain contours to align each brain to the map (Fig. 2D, E). We then compiled lesion outlines from 14 non-seizing and 12 seizing animals into a heatmap (Fig. 2F), highlighting affected areas in seizing and non-seizing animals. The image suggests that larger lesions (manifested as the warm-colored margins) and more anterior located lesions (manifested as the orange/red frontal area) are associated with seizing phenotype more often.

We further stratified lesions by cortical functional area (Fig. 2G). Due to high inter-animal variability in lesion size and location, this analysis remains descriptive. To improve interpretability, we consolidated cortical regions into three functional domains: *Sensory*, *Executive*, and *Limbic/Associative*. These reflect dominant connectivity patterns—input-dominated, output-dominated, or extensively interconnected with subcortical structures. Lesions in seizing animals more frequently involved the *Executive* areas than those in non-seizing animals (Fig. 2H).

We compared lesion sizes between groups and found no significant difference: 10.2 (6.9–14.8) mm² in seizing vs. 6.8 (5.3–10.9) mm² in non-seizing animals (Mann–Whitney *U*-test, *p* = 0.43, Fig. 2I). However, the frontal position of the lesion related to bregma proved to be a significant indicator corresponding to epileptogenicity, 7.4 (6.5–7.8) mm in seizing mice compared to 6.1 (5.6–6.4) mm in non-seizing mice (Mann–Whitney *U*-test, *p* = 0.007, Fig. 2J). Although lesion size did not differ significantly between seizing and non-seizing animals in our dataset, it is often hypothesized to contribute to seizure susceptibility. We therefore propose a composite “lesion severity index” (LSI) that integrates both lesion size and position (Fig. 2K). LSI was also a significant factor of seizure development (Mann–Whitney *U*-test, *p* = 0.005), with seizing mice exhibiting a higher LSI than non-seizing mice, 10.2 (9.7–11.0) in seizing vs. 8.7 (7.8–9.5) in non-seizing animals.

After identifying a significant difference in the rostral–caudal position of the lesion between seizing and non-seizing animals, we asked whether this single spatial parameter could serve as a predictor of seizure occurrence. To test this, we modelled seizure outcome as a function of the anterior–posterior coordinate of the lesion’s frontal edge using logistic regression (Fig. 2L). The model was significant (likelihood ratio test, *p* = 0.0003) with a regression coefficient β₁ = 2.31 (*p* = 0.013). For each animal, we computed a seizure probability based on lesion position. Using a threshold of *p* > 0.5 for positive prediction, the model classified seizure status with over 85% accuracy.

### Cortical mantle in the lesion area is significantly thickened in FCD mice, with prominent effect in seizing animals

Alteration in thickness of the cortical mantle is one of the reported abnormalities in FCD type II. We performed morphological analysis in coronal slices originating from the central part of the lesion. In each mouse, we measured cortical thickness in the center of the lesion and normalized it to the thickness of the homotopic cortex area within the same slice (Fig. 3A). We found that cortical thickening was significantly higher in the FCD animals (n = 16), 9.0 (5.5–19.5) % compared to control animals (n = 8), 1.5 (0–4.5) % (Mann–Whitney *U*-test, *p* = 0.008, Fig. 3D). Furthermore, cortical thickening was significantly more pronounced in seizing FCD animals (n = 7) compared to their non-seizing counterparts (n = 9): 21.0 (11.3–26.3) % vs. 8.0 (4.3–9.8) % (Mann–Whitney *U*-test, *p* = 0.02, Fig. 3G).

**Fig. 3.**
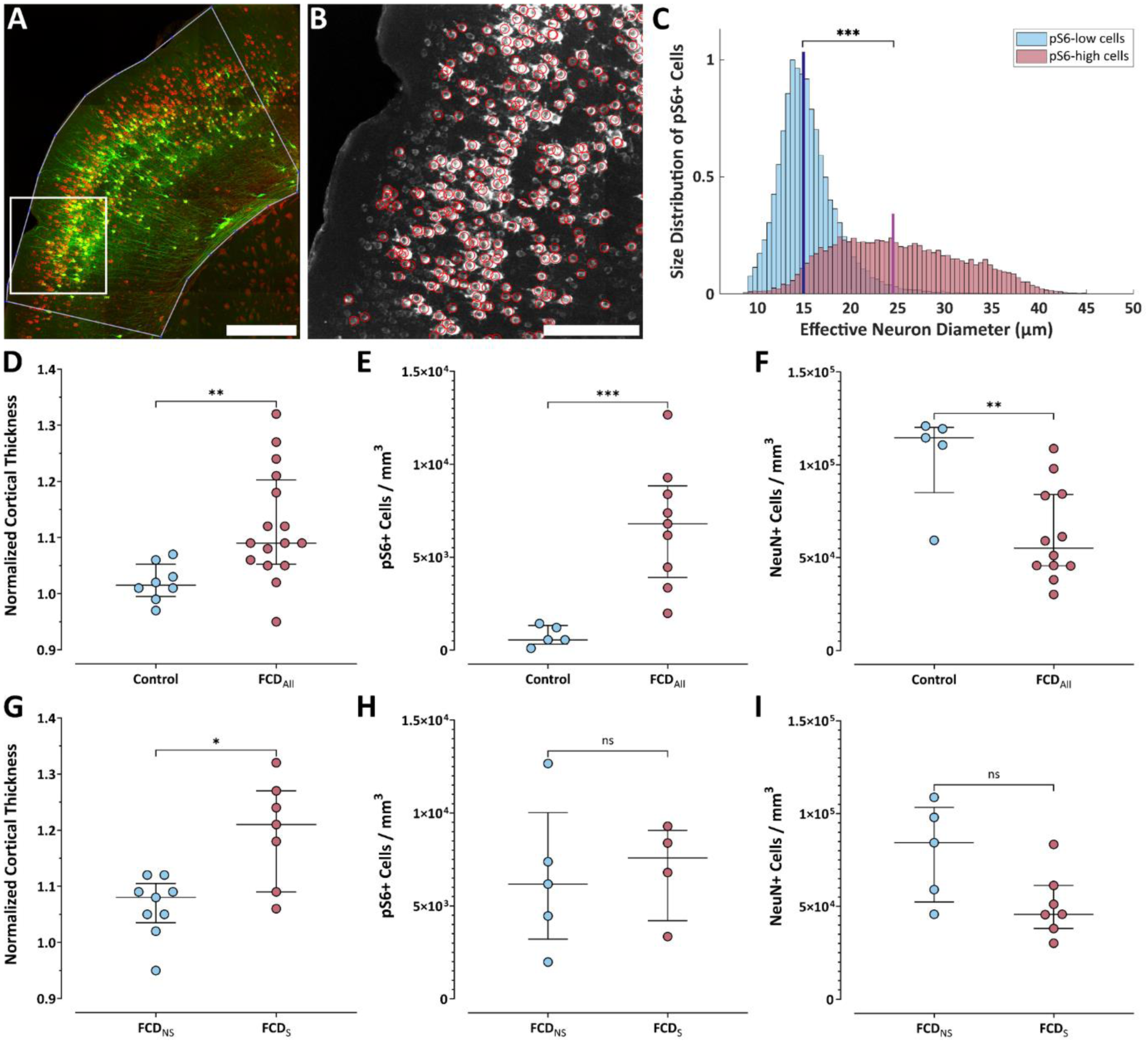
Cortical thickness and cellular properties in FCD. (**A**) Tiled image of coronal brain slice from 2-photon Z-stack maximum projections from the lesion area showing IUE EGFP neurons (green) and pS6 immunostaining (red), with cortical margins outlined; scale bar = 500 μm. (**B**) A cropped section of the red channel taken from the tile inset in (**A**), with red circles indicating pS6+ cells detected by Cellpose that passed intensity thresholding; scale bar = 200 μm. (**C**) Histograms of effective cell diameters of pS6+ cells with either a high (red) or low (blue) pS6 signal (see **Suppl.** Fig. 2). (**D, G**) Cortical thickening in the lesion center normalized to the contralateral hemisphere. (**E, H**) Spatial density of pS6+ neurons. (**F, I**) Spatial density of NeuN+ cells (neurons). (**D–I**) Graphs show comparisons either between all FCD (FCD_All_) and control animals (upper panels), or between FCD animals with (FCD_S_) and without (FCD_NS_) seizures (lower panels); points represent individual animals, central lines = medians, whiskers = IQRs.

### FCD lesions in seizing and non-seizing animals do not differ in density of mTORC1-hyperactive neurons

Phosphorylated ribosomal protein S6 (pS6) is commonly used as a readout of mTOR complex 1 (mTORC1) activity. Traditional assessments often rely on qualitative comparisons or overall staining intensity [11]. To directly examine the relationship between density of mTORC1-hyperactive neurons and seizure susceptibility, we quantified the density of neurons with abnormally elevated mTORC1 activity within FCD lesions.

We performed pS6 immunohistochemistry on brain coronal sections. GFP fluorescence was used to define the lesion area (Fig. 3A). As a novel approach, we developed an algorithm for setting the proper intensity threshold to distinguish neurons with baseline pS6 expression from those with highly increased mTOR activity (Fig. 3B, Suppl. Fig. 2). The algorithm assumes a presence of two neuronal populations with substantially different pS6 levels in the lesion (see *Materials and Methods*). Neurons in FCD that were classified as mTOR-hyperactive based on their pS6 labeling intensity were substantially larger than weakly pS6-labeled neurons (Fig. 3C). Their median diameter was 24.4 (19.6–30.1) μm, which was significantly larger compared to baseline-bright neurons 14.9 (13.3–16.9) μm, (Mann–Whitney *U*-test, *p* < 0.001). As expected, our results revealed a significantly higher density of pS6-positive neurons per mm^3^ in FCD mice (n = 9), 6.8 (4.2–8.6) ×10^3^ cells/mm^3^ compared to control animals (n = 5), 0.6 (0.4–1.3) ×10^3^ cells/mm^3^ (Mann–Whitney *U*-test, *p* = 0.002, Fig. 3E). However, we did not observe any significant difference between the groups of seizing (n = 4) and non-seizing (n = 5) animals; 6.2 (5.1– 8.8) ×10^3^ cells/mm^3^ and 7.6 (3.8–8.7) ×10^3^ cells/mm^3^, respectively (Mann–Whitney *U*-test, *p* = 0.73; Fig. 3H).

### Spatial density of neurons is significantly reduced in FCD lesions compared to control cortex

To assess overall neuronal density within FCD lesions, we quantified NeuN-positive cells in coronal brain sections from FCD and control mice. Our analysis revealed a significant reduction in NeuN-positive cell density within the FCD lesions (n = 12), 5.5 (4.6–8.4) ×10^4^ cells/mm^3^ compared to controls (n = 5), 11.5 (9.8–12.0) ×10^4^ cells/mm^3^ (Mann–Whitney *U*-test, *p* = 0.012, Fig. 3F). We did not observe a significant difference between the seizing (n = 7) and non-seizing (n = 5) FCD mice: 4.6 (4.0–5.9) ×10^4^ cells/mm^3^ vs. 8.4 (5.6–10.1) ×10^4^ cells/mm^3^ (Mann– Whitney *U*-test, *p* = 0.15, Fig. 3I).

Finally, we examined whether seizure development was associated with a higher fraction of pS6+ neurons among all NeuN-positive neurons. In animals for which both pS6 and NeuN immunostaining were available, we calculated the fraction and compared values between seizing and non-seizing animals (Suppl. Fig. 3). We found no significant difference between seizing (n = 4) and non-seizing (n = 5) FCD mice: 12.6 % (9.4–17.8) vs. 7.3 % (3.9–17.4) (Mann–Whitney U-test, p = 0.56).

### Altered axonal and synaptic bouton properties in FCD type II

We analyzed axonal properties in mounted brain slices from FCD and control animals using confocal microscopy (Fig. 4A). High-resolution scans were acquired from defined regions of interest (ROIs) on both the ipsilateral side (Fig. 4B) and the contralateral projections (Fig. 4C), allowing us to resolve individual axons in all examined regions. Imaging was conducted at three cortical depths that correspond to layer 2/3 (L2/3), layer 5 (L5), and the border between layer 6 and the white matter (WM). Axonal morphology was assessed on both sides, and individual axons were reconstructed in 3D (Fig. 4 D-F). Axonal diameters were measured along their trajectory (Fig. 4G) to quantify shaft thickness and to detect axonal boutons (Suppl. Fig. 4). Bouton diameter, volume, linear density, and axonal shaft thickness were compared using the Mann–Whitney *U*-test.

**Fig. 4.**
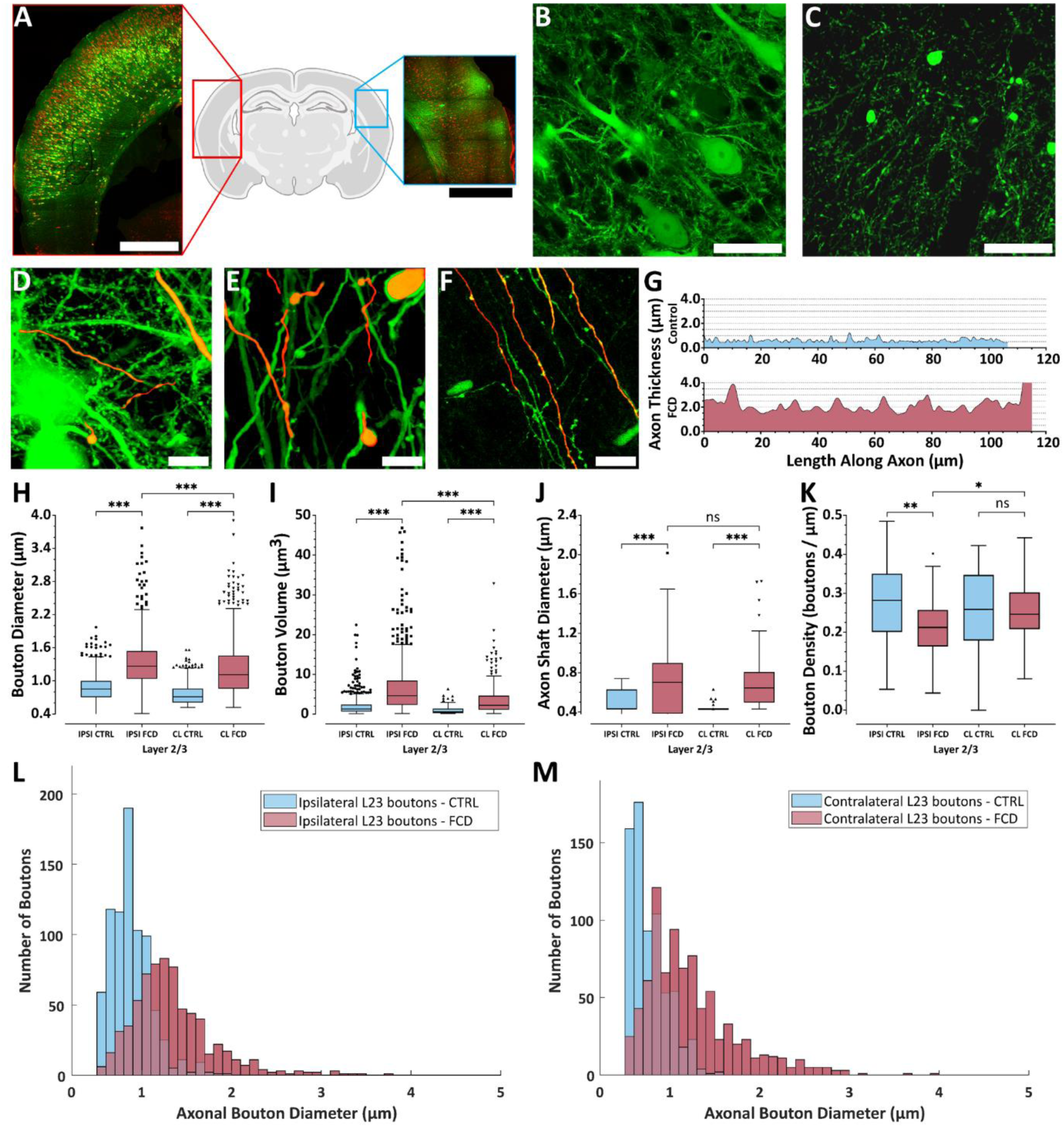
Axon shafts and boutons are enlarged in FCD, but bouton density is reduced. (**A**) Tiled two-photon maximum projection of a coronal brain slice showing the typical appearance of ipsilateral (red) and homotopic contralateral (blue) regions; scale bars = 1 mm. (**B**) Confocal image of GFP+ pyramidal neurons in ipsilateral layer 2/3; scale bar = 100 μm. (**C**) Confocal image of GFP+ axonal projections in the homotopic contralateral layer 2/3; scale bar = 100 μm. (**D-F**) Traced axons (red) from ipsilateral layer 2/3 (**D**), and layer 5 (**E**) in an FCD animal and in ipsilateral layer 2/3 in a control FCD animal (**F**); scale bars = 20 μm. (**G**) Quantification of axon shaft diameters in layer 2/3. (**H–K**) Tukey boxplots of bouton diameter (**H**), bouton volume (**I**), axon shaft diameter (**J**), and bouton linear density (**K**), measured in layer 2/3. (**L, M**) Histograms of bouton diameters (**H)** in layer 2/3 from control and FCD animals on the ipsilateral (**L**) and homotopic contralateral sides (**M**). (**H–M**) Data from layer 2/3. CTRL = control animals; FCD = lesioned animals; IPSI = ipsilateral side; CL = contralateral side.

The mTOR pathway, *inter alia*, critically regulates synaptic bouton growth through control of protein synthesis and cytoskeletal dynamics [29]. Recently, abnormally large axonal varicosities—presumed to be boutons—have been reported in contralateral projections of dysmorphic neurons[16]. Here, we provide a quantitative analysis of axonal shaft thickness and bouton morphology, including their diameters, volumes, and linear densities. These parameters were evaluated in six regions across both hemispheres: L2/3 (Fig. 4H–M), L5, and WM (Suppl. Fig. 6). Analysis revealed that boutons in FCD mice were significantly larger in diameter compared to those in control mice across all areas (L2/3, L5, and WM) on both the ipsilateral and contralateral sides (Fig. 4H). In L2/3, the median bouton diameters for ipsilateral projections were 0.85 (0.71–1.00) μm in control animals and 1.30 (1.00–1.50) μm in FCD animals. For contralateral projections, the respective median bouton diameters were 0.71 (0.61–0.86) μm in control animals and 1.10 (0.86–1.50) μm in FCD animals. Notably, boutons within the FCD lesion itself were even larger than those observed in contralateral projections. All comparisons were statistically significant (Mann–Whitney U-test, *p* < 0.001).

Similar and highly significant differences were also observed in bouton volume measurements (Fig. 4I; Table 2). Axonal shafts were significantly thicker in FCD animals compared to controls on both the ipsilateral and contralateral sides (Fig. 4J). On the ipsilateral side, median shaft thicknesses were 0.70 (0.38–0.90) μm in FCD axons and 0.43 (0.43–0.63) μm in controls. Comparable results were found contralaterally: 0.63 (0.50–0.80) μm in FCD axons versus 0.43 (0.43–0.43) μm in controls. All comparisons were statistically significant (*p* < 0.001). Within each group (FCD or control), no significant differences in shaft thickness were found between hemispheres.

**Table 2.**
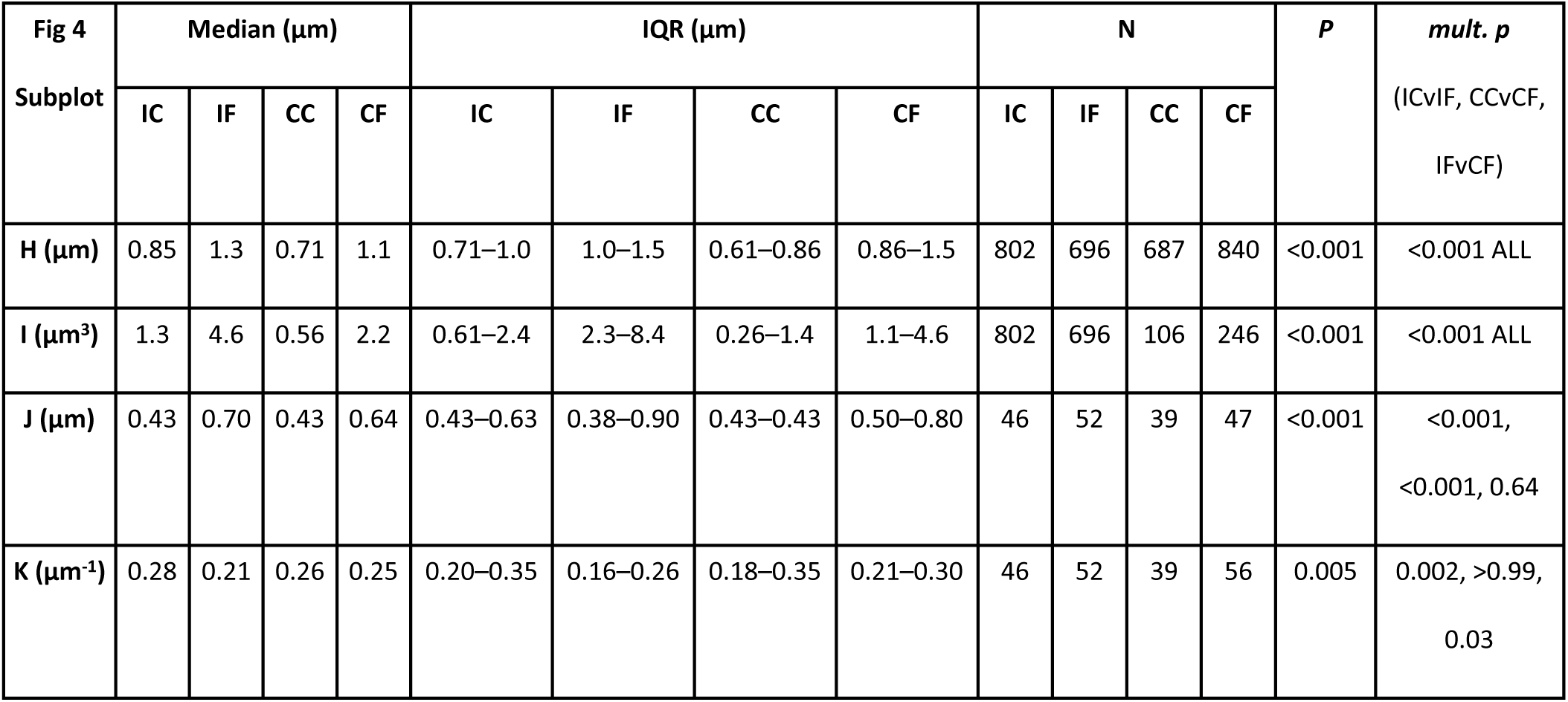
Complete values to Fig. 4. Legend: IC = ipsilateral control; IF = ipsilateral FCD; CC = contralateral control; CF = contralateral FCD; IQR = interquartile interval; N = number of boutons/shafts, mult. *p* = *p* values from multiple comparisons tests.

Surprisingly, bouton density did not increase in FCD axons. On the ipsilateral side, FCD axons exhibited 0.21 (0.16–0.26) boutons/μm, significantly lower than the 0.28 (0.20–0.35) boutons/μm observed in controls (Fig. 4K).

Of particular note, massively enlarged boutons were observed in FCD, far exceeding the diameter and volume typical for presynaptic terminals, making median values alone insufficient to describe the data. To better illustrate these outliers, we plotted diameter and volume distributions for both hemispheres (Fig. 4L, M). Analogous findings were observed in L5 and WM (Suppl. Fig. 5 and Suppl. Table 1), with one exception: bouton density in FCD axons, where no significant side-to-side difference emerged.

### Axonal varicosities (boutons) were positive for synapse-associated proteins in the contralateral cortex, but rarely in the corpus callosum

To assess the synaptic nature of giant axonal boutons formed by dysmorphic neurons, we performed immunohistochemistry using antibodies against vGlut1 and Synapsin-1. Punctate labeling for both proteins was frequently observed within boutons located on axons in the contralateral cortex, but only rarely within boutons on axons traversing the corpus callosum (Suppl. Fig. 6A, B and D, E, respectively).

We further validated these findings in brain slices from three mice that had been electroporated with a different plasmid mixture, in which dysmorphic neurons expressed Cre recombinase and the infrared fluorescent protein iRFP_713_ instead of EGFP (see *Materials and Methods* for details on the co-electroporation plasmid). These neurons were transduced with an AAV vector to Cre-dependently express soluble tdTomato and a synaptophysin-EGFP fusion protein. In this system, synaptophysin-EGFP clusters were readily detected within red-filled boutons, facilitating colocalization by restricting fluorescence exclusively to the axons and boutons of transduced neurons (Suppl. Fig. 6C, F).

Finally, we stained for PSD95 to assess postsynaptic contacts at these excitatory boutons. Despite their large size, the boutons were not associated with an increased number of apposed PSD95-positive puncta that would suggest multiple synaptic connections. On the contrary, our data indicate that even giant boutons typically form only a single postsynaptic contact (Suppl. Fig. 6G).

### Electroporation of two plasmids led to high overlap of pS6+ and EGFP+ neurons in our FCD model

We co-electroporated a second plasmid encoding EGFP under the strong CAG promoter, enabling robust and consistent labeling of transfected neurons. To verify that the vast majority of EGFP-positive neurons also expressed the mutated *mTOR* gene construct, we assessed the overlap between EGFP fluorescence and mTOR pathway hyperactivation, as indicated by pS6 immunostaining. A background-independent classification algorithm was used to distinguish neurons transfected with the *mTOR* construct from non-transfected cells, based on pS6 signal intensity. In control animals, pS6 staining was detectable in a subset of neurons (Fig. 5A), but its intensity was substantially lower than in the strongly labeled, cytomegalic neurons observed in FCD animals (Fig. 5B). To directly quantify the overlap between EGFP expression (from the second plasmid) and mTOR hyperactivation (from the large *mTOR* plasmid), we thresholded the pS6 immunofluorescence and calculated the proportion of EGFP+ neurons that also showed high pS6 signal. This analysis revealed that a median of 75% (63–81%) of EGFP+ neurons in FCD animals were also pS6-high (Fig. 5C, Mann–Whitney *U*-test, *p* < 0.001), indicating a strong correspondence between EGFP labeling and mTOR pathway activation. In contrast, only 0–3% of EGFP+ neurons in control animals showed comparable pS6 signal. The size distribution of EGFP-positive neurons further supported this distinction: neurons within FCD lesions were significantly larger than those in the corresponding regions of control animals (median diameter: 27.0 [23.4–30.6] µm vs. 15.7 [14.4–17.4] µm, Mann–Whitney *U*-test, *p* < 0.001; Fig. 5D). We therefore treated GFP+ cells as a proxy for mTOR-mutated neurons in subsequent analyses.

**Fig. 5.**
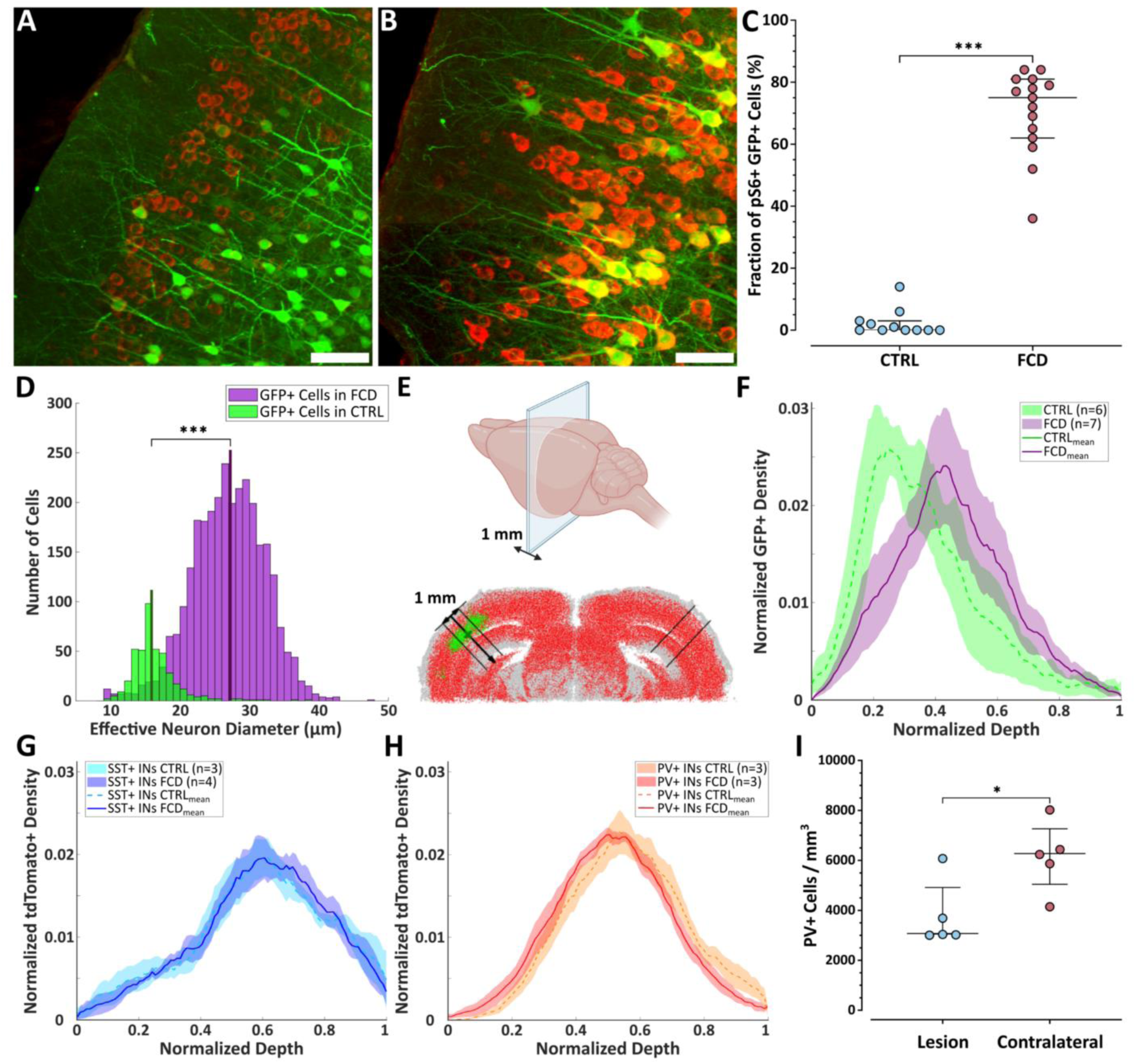
Cell type-specific quantification using light-sheet microscopy or classical immunostaining. (**A, B**) Two-photon Z-stack maximal projections of coronal brain slices acquired in the ipsilateral lesion area of a control animal (**A**) and an FCD animal (**B**), stained for pS6 (red) and showing EGFP expression (green); scale bars = 100 μm. (**C**) Percentage of EGFP+ neurons classified as pS6-high in FCD and control brains; points represent individual animals, central lines = medians, whiskers = IQRs. (**D**) Histogram of effective diameters of EGFP+ neurons in FCD (green) and control (purple) animals. (**E**) Schematic of light-sheet–based cell density calculation along the cortical depth axis. (**F**) Cortical depth profile of EGFP+ neuron distribution in FCD lesions and control samples. (**G, H**) Same analysis as in (**F**), shown for SST+ interneurons (**G**) and PV+ interneurons (**H**). (**I**) Spatial densities of PV-immunolabeled neurons in FCD animals (n = 5), comparing ipsilateral and contralateral cortical regions; points represent individual animals, central lines = medians, whiskers = IQRs.

Using light-sheet microscopy, we evaluated the depth distribution of EGFP-positive neurons in the core of the lesion in cleared brains (Fig. 5E). Although the group-averaged curves appeared different (Fig. 5F), principal component analysis (PCA) of curve shapes did not demonstrate a significant group difference (PC1 p = 0.09).

### Unchanged distribution of PV+ and SST+ interneurons in FCD, with reduced PV+ density

Using light-sheet microscopy, we analyzed the depth distribution of somatostatin-positive (SST+) and parvalbumin-positive (PV+) interneurons in FCD and control brains of SST-IRES-Cre/flex-tdTomato and PV-P2A-Cre/flex-tdTomato crosses. No significant differences in the shape of the laminar distribution were observed for either SST+ (Fig. 5G) or PV+ interneurons (Fig. 5H). The group-averaged curves nearly overlapped across both cell types and both conditions, and PCA confirmed the absence of group differences (PC1: p = 0.24 for SST+, p = 0.95 for PV+). Based on previous reports of reduced PV+ interneuron density in FCD [27, 30, 31], we aimed to validate this observation in our FCD model using conventional PV immunohistochemistry (Fig. 5I). To control for seizure-related variability and regional differences, we compared PV+ cell densities between the FCD lesion and the homotopic contralateral cortex within the same brain slices from five seizing C57BL/6J mice with FCD. This analysis revealed a significant reduction (up to 50%) in PV+ neuron density within the FCD lesion: 3.1 (3.1– 4.3) × 10³ cells/mm³ compared to 6.3 (5.5–6.9) × 10³ cells/mm³ in the contralateral cortex (Mann–Whitney *U*-test, *p* = 0.03).

## DISCUSSION

### The *in utero* mouse model of FCD type II recapitulates key histological and electrophysiological features

To study FCD in mice, we employed the FCD type II model introduced by Lim et al. [11], which utilizes the patient-derived mutation p.L2427P. The model recapitulates key histopathological features: the presence of DNs, cortical thickening, white matter heterotopia, and loss of cortical lamination [30–32]. Consistent with most FCD mouse models [11, 27, 30, 33, 34], balloon cells (BCs), a hallmark of FCD type IIb in humans, were not observed. The absence of BCs in rodent FCD II models likely reflects both species-specific and developmental differences and probably cannot be explained by IUE timing alone. The human cortex contains an expanded outer subventricular zone rich in outer radial glia, structures that are sparse in rodents and considered critical for generating BC-like phenotypes [35, 36]. Regarding astrocytic origin, the role of IUE timing can be excluded based on experiments in which *TSC1/TSC2* were knocked down using CRISPR/Cas9 at E14 in radial glial progenitors— expected to give rise to mutated astrocytes—but no BCs were observed [37]. BC-like “giant cells” appeared only in a stronger and earlier *Tsc1* loss model, showing mixed glial/neuronal identity and mitochondrial defects closely resembling human pathology [38]. Altogether, these findings suggest that BC formation depends on both species- and development-specific context together with the timing and strength of the mutation. We did not perform any immunohistochemical staining for markers typical for balloon cells in humans, such as GFAP or vimentin.

We confirmed that the mice developed spontaneous seizures; however, these occurred in less than half of the animals. This proportion is generally lower than that reported previously in FCD animal models (see below). Such discrepancies may arise from differences in the specific genetic mutations used, plasmid concentrations, localization of the lesion, and the genetic background of the mouse strain.

Several different genetic variants in PI3K/AKT/mTOR pathway have been identified in FCD type II patients and evaluated in animals. For instance, introducing the *RHEB* p.P37L mutation [16], which renders the RHEB protein out of the control of TSC1/TSC2 complex, and targeting the somatosensory cortex (SSC) in FvB/NHsD mice reportedly resulted in seizures in all animals, while the *RHEB* p.S68P variant [39] induced seizures in 83% (5/6) in similarly prepared animals. With constitutively active *RHEB* p.S16H variant [27], which is hyperactive but remains responsive to TSC1/TSC2 regulation, spontaneous seizures occurred in 73% (8/11) of animals with the lesion in the medial prefrontal cortex (mPFC), specifically the anterior cingulate cortex (ACC). However, no seizures were observed when the same construct targeted the SSC. In another study, using *RHEB* p.S16H variant to induce FCD in the mPFC of CD1 mice, seizure incidence depended on the plasmid concentration and reached 100% at higher concentrations [28]. A recent study employed IUE of a construct for CRISPR/Cas9 components to knock out *DEPDC5*, an inhibitor of the amino acid-sensing branch of the PI3K/AKT/mTORC1 pathway, in the motor cortex [34]. This led to seizures in only 30% (4/13) of Swiss-Webster mice.

In our study, 45% of the animals with FCD developed seizures (Fig. 1), which is in contrast with 90% seizure incidence reported by Lim et al. [11], despite using the same mutation and the same mouse strain (C57BL/6J). We hypothesize that variability in lesion size and location may explain this discrepancy between otherwise identical models. Apart from seizures, the mouse model of FCD used in this study exhibits rich interictal activity [26]. We found that FCD mice without seizures showed a low spike rate, slightly but not significantly higher than in non-epileptic controls, consistent with our previous observation that the increase in interictal activity mostly follows the onset of seizures [40].

### Lesions in frontal cortex promote seizure occurrence in the FCD model

In animal models, lesion placement is often underreported. We therefore evaluated how lesion size and location influence seizure propensity in the mTOR p.L2427P mutation FCD model. In our cohort, there was no statistically significant difference in lesion size between animals that developed seizures and those that did not. In contrast, the position of the lesion proved to be a significant factor. Our data indicate that seizures are more likely to develop when the lesion is located in the frontal cortex, particularly in those involving the primary motor, secondary motor, and frontal association areas (Fig. 2). Logistic regression analysis confirmed that the position of the lesion is a meaningful predictor of seizure occurrence. An appealing hypothesis is that the portion of the lesion affecting frontal cortical regions may play a decisive role in the generation and maintenance of epilepsy. If this hypothesis is correct, a partial resection confined to the frontal part of the lesion might provide seizure control comparable to complete resection, while reducing the risk of adverse functional outcomes.

In general, our observations are consistent with clinical studies [5, 41, 42], which report that FCD type II lesions are most frequently identified in extratemporal regions, particularly in the frontal cortex. Our data raise the possibility that a subset of “silent” FCDs exists and that their quiescence may reflect a predominance in parietal or occipital regions. The only systematic study of incidentally detected FCDs [43] reported a frontal predominance (>60%), but this may have been biased by the indications for MRI. Notably, frontal FCDs appear more epileptogenic, whereas temporal and occipital lesions associated with epilepsy tend to be larger [3, 37], potentially compensating for their less favorable location by involving a greater cortical area.

The frontal predominance of FCD is likely multifactorial, involving both network embedding and regional molecular context. Frontal association areas function as hub-rich regions with dense reciprocal connectivity, where even small lesions can recruit large-scale networks and stabilize epileptogenic loops— consistent with human connectome data showing that over half of frontal FCDs overlap multiple association networks [44] and with mesoscale mouse studies revealing strong interconnectivity among prefrontal and cingulate areas [45]. Cell-type–specific tracing further confirms a hierarchical and reciprocally connected architecture that supports hub-like dynamics [46]. Additional factors probably include the late developmental trajectory of frontal cortex, extending vulnerability to somatic mTORopathies [47], and the lower baseline MTOR expression in frontal regions, which may amplify the relative impact of mutations [44]. Finally, single-nucleus data show mitochondrial and synaptic dysregulation in dysmorphic neurons with excitatory changes in neighboring cells, linking local metabolic stress and hub embedding to seizure generation [48].

### Cortical thickening alone does not explain reduced neuronal density in our model

Increased cortical thickness at the lesion site compared to adjacent cortex is a documented morphological abnormality in FCD type II patients [7, 32, 49, 50]. In animal models, two previous studies [27, 33] described the changes in the cortical mantle thickness. Hsieh et al. [27] measured the cortical thickness in a mouse model of FCD using magnetic resonance imaging in 3- to 4-month-old animals and reported an approximately 10% increase at the lesion site relative to the contralateral hemisphere. In our dataset, we observed a comparable median increase of ∼9% across all FCD animals. Notably, this increase was significantly greater (median 21%) in animals that developed spontaneous seizures (Fig. 3). These volumetric changes can likely be attributed, at least in part, to the enlarged somata and dendrite ramification of DNs, a potentially greater contribution of glial cells to the tissue volume [27, 28], and increased extracellular space due to excessive deposition of extracellular matrix components [12]. Determining the relative contribution of each of these factors will require future comprehensive study, performed within the same animal model or, preferably, within the same individual subjects.

NeuN is a widely used marker of mature neurons in both healthy and pathological brain tissue. In human FCD samples and corresponding animal models, reduced NeuN immunoreactivity has been frequently interpreted as indicative of neuronal loss. Within the context of FCD, the interpretation of reported reductions in neuronal density remains controversial. In human patients, some studies report very large reduction in NeuN-positive cell density within FCD lesions, with decreases exceeding 50% [41], and in some cases further declining with the duration of epilepsy [51]. However, other research groups challenge this view by reporting no significant difference in neuronal density compared to controls, even in FCD IIB patients [52]. In animal models, a recent study using an IUE-based model of FCD II reported a 20–25% decrease in neuronal density [33] that could be attributed to tissue expansion (effective increase of the denominator of density calculations). In our dataset, we observed a significant and substantial reduction in NeuN-positive cell density - 50% - in FCD tissue compared to controls (Fig. 3). This decrease was present regardless of seizure occurrence. If the observed ∼9% increase in cortical thickness reflected isotropic expansion (i.e., 9% increase in all three dimensions), it would correspond to an overall volumetric increase of approximately 30%, which alone cannot explain the magnitude of the observed decrease in neuronal density. Because we did not observe any EGFP-positive cellular debris or remnants in any of the FCD sections, we consider overt neuronal loss due to cell death an unlikely explanation in our model. Alternatively, the reduced neuronal density may reflect a lower local number of neurons due to disrupted migration and proliferation during cortical development [53], a greater degree of tissue expansion in tangential directions (i.e., perpendicular to cortical depth), or simply the large natural variability in neuronal density across the dorsal cortex—for example, neuron density in the visual cortex can be nearly twice as high as in the motor cortex [54, 55].

### No significant differences in pS6+ neuronal density or fraction in FCD between seizing and non-seizing animals

In this study, primary antibodies against pS6 were used as a proxy for the mTOR pathway activity level. In FCD, we observed two neuronal populations that were highly separable based on their intensity of pS6 labeling. We developed a simple algorithm for identification and separation of these two groups (Suppl. Fig. 2). This method offers a reproducible metric that could aid future studies in establishing a threshold for pathological mTOR activation solely based on one’s acquired data.

Our data did not show significant differences in pS6+ neurons’ spatial density between the FCD samples from seizing animals compared to non-seizing animals. However, this could be caused by the large variance in different volumetric changes in individual FCD mice and the natural neuronal density of the cortex in which the respective neuron resided. To partially remove that variability, we calculated the fractions of pS6+ spatial density and NeuN-positive cell density in consecutive slices from the same individual mice (Suppl. Fig. 3). Although our results indicate that the median fraction of pS6+ neurons out of all neurons (∼13 %) in FCD of seizing animals is almost double to the value in non-seizing animals (∼7%), our results are not significant due to still large variability and low sample size. The variability could potentially be caused by the fact that the density values of pS6 and NeuN were not from identical slices. We hypothesize, that a careful experiment with double immunolabeling within the identical slices and a big cohort of animals will be necessary for resolving the question. Similar results were indicated by other authors in the form of representative images, although the load was not quantified in terms of ps6+ neuron and NeuN spatial density [11, 27, 28, 56]. In one influential study [28], the authors showed that the epilepsy developed only in animals electroporated with intermediate concentration of plasmid that carried a mutated gene from the mTOR pathway with higher seizure frequency in animals with highest plasmid concentration. On the other hand, they showed, that although the morphological changes of the transfected neurons, such as size, were altered proportionally to the used plasmid concentration, the density of transduced neurons in the cortex was not different between the three plasmid concentrations. Another alternative, partially mimicked by the methodology in [28], is that the type of the mutation and its influence on the overall activity on mTOR complexes governs the epileptogenicity of the lesion regardless of the mutated neuron fraction. Those neurons can then influence other neurons by non-cell autonomous effects [57, 58], which can thus spread the alterations to other neurons, hypothetically in a mutation severity-dependent manner.

In human epilepsy patients with FCD type II with identified causal mutation in the mTOR pathway, the frequency of the alternative allele has been repeatedly reported as typically between 1–15% [7, 59, 60]. Across all our mice with FCD, our reported value of (10%) lies within this range.

### Fluorescent marker from the main plasmid carrying the mutated mTOR was insufficient for lesion visualization and axonal morphology analysis

The main plasmid carrying the mutated *mTOR* gene was relatively large (13.9 kbp, with *mTOR* cDNA spanning 7.6 kbp), which likely reduced IUE transfection efficiency. Its fluorescent marker, mAmetrine, was expressed as a second gene in a bicistronic cassette via an IRES linker, further limiting its expression. As a result, identifying transfected cells or selecting pups with visible lesions was unreliable, and the signal was too weak to delineate lesions through the skull or on explanted brains. Moreover, the fluorescence intensity was insufficient for high-resolution imaging of fine neuronal structures, such as axonal shafts and individual boutons. While the lower transfection or expression efficiency may have limited fluorescent protein detection, the mutated mTOR was still sufficiently potent to induce the FCD phenotype. To address the insufficient fluorescence, we co-electroporated a second, smaller plasmid expressing EGFP under a strong promoter (Fig. 1B). Colocalization of GFP and pS6 immunoreactivity showed that up to >75% of GFP+ cells in FCD lesions had high mTORC1 activity (Fig. 5C).

### Dysmorphic neuron axons and boutons show abnormal morphology across both hemispheres

Axonal abnormalities have previously been documented in FCD type II, although their functional consequences remain incompletely understood. *In vitro* experiments using overexpression of the *RHEB* p.P37L gene mutation in primary hippocampal neuron cultures demonstrated that hyperactivation of the mTOR pathway causes increased axonal length and branching [16]. Similarly, loss of TSC1 or TSC2 function—endogenous inhibitors of the mTOR pathway—induced ectopic axonal growth in the mouse cortex [61]. In a mouse model of FCD type II, RHEB p.P37L overexpression led to a broader distribution of contralateral axonal terminals, extending into multiple layers of homotopic and neighboring cortical areas [16].

We analyzed axonal morphology and synaptic bouton properties in a mouse model of FCD type II, both contralaterally and ipsilaterally within the lesion core. In L2/3, axonal shafts in FCD mice were approximately twice as thick as in controls on both sides of the cortex (Fig. 4J); similar differences were observed in L5 and WM (Suppl. Fig. 5C, G). Given our use of high-resolution confocal microscopy and cytoplasmic GFP, we interpret this as an increased axoplasmic cross-section. Myelin thickness was not assessed. There are only limited data describing myelination changes in FCD. One group [62] reported minor but statistically significant differences in myelination between FCD type IIA and IIB human samples based on histopathological analysis. Their findings were mixed: medium-caliber axons in FCD IIA showed slightly reduced myelin sheath thickness, whereas large-caliber axons in FCD IIB exhibited increased thickness. However, it is not entirely clear how axons of mutated neurons were identified in the electron microscopy preparations, or how the reported structural changes relate to the unaltered levels of myelin-related mRNA observed in FCD IIA neurons. Overall, this area of research requires careful reassessment before firm conclusions can be drawn about the extent and functional relevance of myelination changes in FCD type II.

We found giant synaptic boutons on both sides of the brain. On the ipsilateral side, high-resolution Z-stacks from confocal microscopy revealed boutons up to 5 μm in L2/3, with median diameters and volumes 1.5× and 3.4× larger than in controls (Fig. 4). Across all analyzed regions, bouton sizes were consistently larger within the FCD lesion.

Onori et al. [16] interpreted giant boutons as glutamatergic synaptic terminals based on vGlut and Synapsin immunostaining and a representative EM image showing synaptic vesicles in a single bouton from a mutated neuron, alongside one from a control neuron. We also investigated the nature of these structures. Slices were immunostained for vGlut1, Synapsin1, and PSD95, and we additionally used synaptophysin-EGFP fusion protein expressed in dysmorphic neurons. This fusion marker offered excellent contrast and allowed unambiguous attribution of fluorescent puncta to individual axons. We examined axonal varicosities in the corpus callosum (CC) and in the contralateral cortex homotopic to the FCD lesion. While contralateral boutons consistently colocalized with presynaptic markers, those in the CC largely did not. This suggests that bouton swelling may not be caused by vesicle accumulation and that DNs likely do not form functional synapses in the CC. To assess whether large boutons in the contralateral cortex enhance output via multiple synapses, we stained for PSD95 and found only 1–2 puncta per bouton, indicating normal postsynaptic targeting.

While previous studies demonstrated axonal sprouting of DNs, we found that their axons exhibit a reduced linear density of boutons compared to controls. This suggests that despite increased axonal arborization and bouton size, the overall synaptic output of dysmorphic neurons may be limited.

From a different perspective, the giant boutons may reflect a compensatory response to impaired synaptic machinery. Several studies in rodent epilepsy and dysplastic cortex models report loss of synaptic vesicle glycoprotein 2A (SV2A), indicating synapse reduction and altered neurotransmission [63–65]. This reduction has also been confirmed in FCD type II patients, specifically in the epileptogenic zone [66]. Another open question is whether these enlarged boutons may mediate non-synaptic effects of dysmorphic neurons, including senescence-associated secretory activity [60] such as interleukin-1β or TNF-α release [67].

### PV+ and SST+ interneurons in FCD using light-sheet microscopy and IHC

Alterations in PV+ interneurons in FCD have been inconsistently reported. While some human studies describe a reduction in their number or proportion [31, 52, 68], others report no change [69] or even an increase [70]. Changes in spatial distribution without altered overall numbers have also been described [69, 71]. Most animal models show no significant difference in PV+ interneuron numbers [27, 58], with one study suggesting that apparent decreases may reflect tissue expansion rather than true cell loss [33]. SST+ interneurons are generally considered more resistant to pathological conditions such as oxidative stress and excitotoxicity. Nonetheless, some studies [33, 58] report reduced SST+ numbers in FCD using mRNA in situ hybridization method, particularly in seizure-prone areas.

Using light-sheet microscopy, we found no differences in the depth distribution of PV+ or SST+ interneurons in FCD compared to controls. The CUBIC protocol moderately reduced tdTomato fluorescence and lowered the signal-to-noise ratio, which limited the reliability of automated detection and increased the risk of false positives and negatives. To address interneuron density directly, we performed classical PV immunostaining in a separate group of FCD animals and compared PV+ neuron density between the lesion and the contralateral cortex within the same slices. This analysis revealed a significant reduction in PV+ neuron density in FCD, paralleling the overall decrease in NeuN+ cell density. The density of PV+ interneurons in the contralateral cortex was consistent with to the values previously reported for healthy mouse cortices [72]. We did not perform mRNA in situ hybridization, and conventional IHC is not suitable for identifying SST-positive somata, as SST is a secreted neuropeptide and therefore scarce in cell bodies.

## CONCLUSION

Our study demonstrates that lesion location, particularly within frontal cortical areas, is a strong predictor of spontaneous seizures in a mouse model of FCD type II, whereas lesion size alone is not. We confirmed key pathological hallmarks, including cortical thickening, neuronal hypertrophy, mTORC1 hyperactivation, and enlarged axonal boutons.

Giant boutons from dysmorphic neurons were detected not only contralaterally, but also within the lesion and in the corpus callosum. Whereas contralateral boutons exhibited synaptic features, callosal varicosities largely lacked presynaptic markers. Despite axonal hypertrophy, bouton density was reduced, leaving the net synaptic output uncertain.

Classical immunostaining revealed a proportional reduction of PV+ interneurons and NeuN+ neurons in the studied FCD model.

Together, these findings refine our understanding of seizure risk in FCD and highlight the contribution of lesion topography and cellular microstructure to epileptogenesis.

## DECLARATIONS

### Availability of data and material

The datasets used and analyzed during the current study available from the corresponding author on reasonable request.

## Authors’ contributions

According to CRediT taxonomy the contributions of individual authors were as follows: Conceptualization: NP, PJ, and ON. Methodology: NP, CVLO, MTR, MR, JK, and ON. Software: CVLO, MTR, and ON. Formal analysis: NP, CVLO, MTR, and ON. Investigation: NP, NS, and ON. Data curation: NP, CVLO, MTR, NS, MR, JK, and ON. Writing: NP, CVLO, JP, PJ, and ON. Visualization: NP, CVLO, MTR, and ON. Supervision: JK, PJ, and ON. Project administration: NP, PJ, and ON. Funding acquisition: PJ and ON.

## Acknowledgements

The authors would like to thank Pavlína Smolková for plasmid preparations (amplification) and Paulína Žideková for her assistance with data transport and management. The authors also thank the EpiReC consortium for their support. Figures 1A, 4A, and 5E were partially created with BioRender.com.

## SUPPLEMENTARY FIGURES

**Supplementary Fig. 1.**
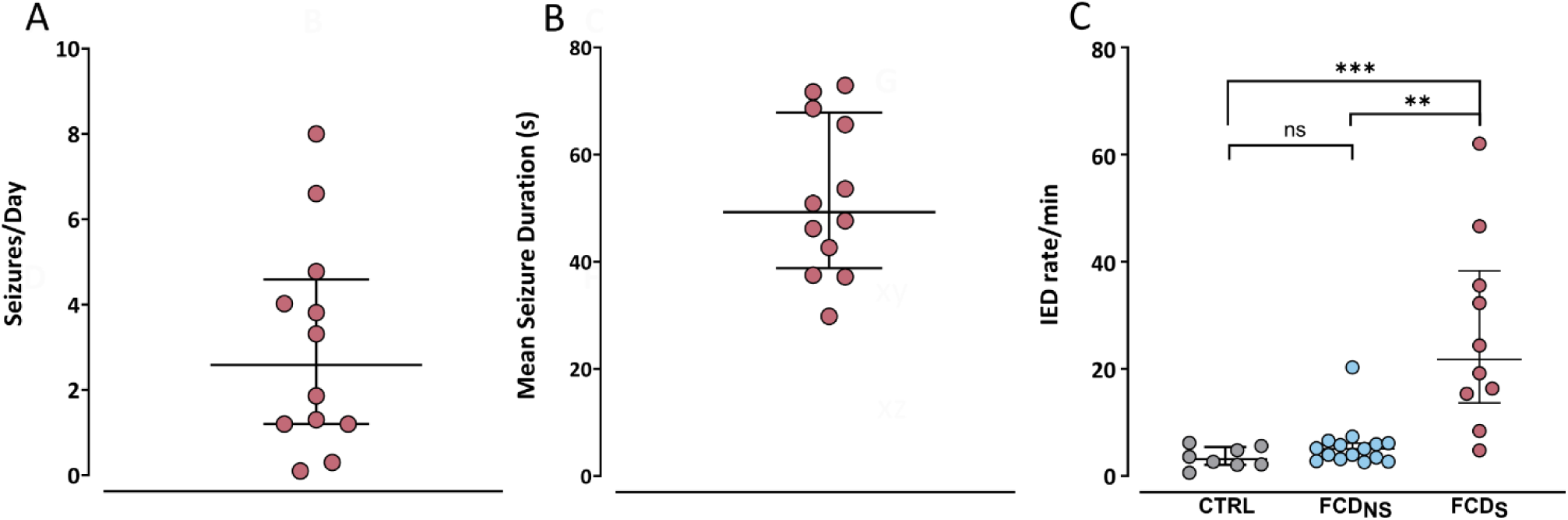
Epileptic phenotype of the animals involved in the study. **(A)** Number of seizures per day and **(B)** their mean duration in all recorded mice; each point represents an individual animal, horizontal lines indicate medians, and whiskers denote interquartile ranges (IQRs). **(C)** Comparison of interictal epileptiform discharge (IED) frequency between control animals and FCD mice without or with seizures.

**Supplementary Fig. 2.**
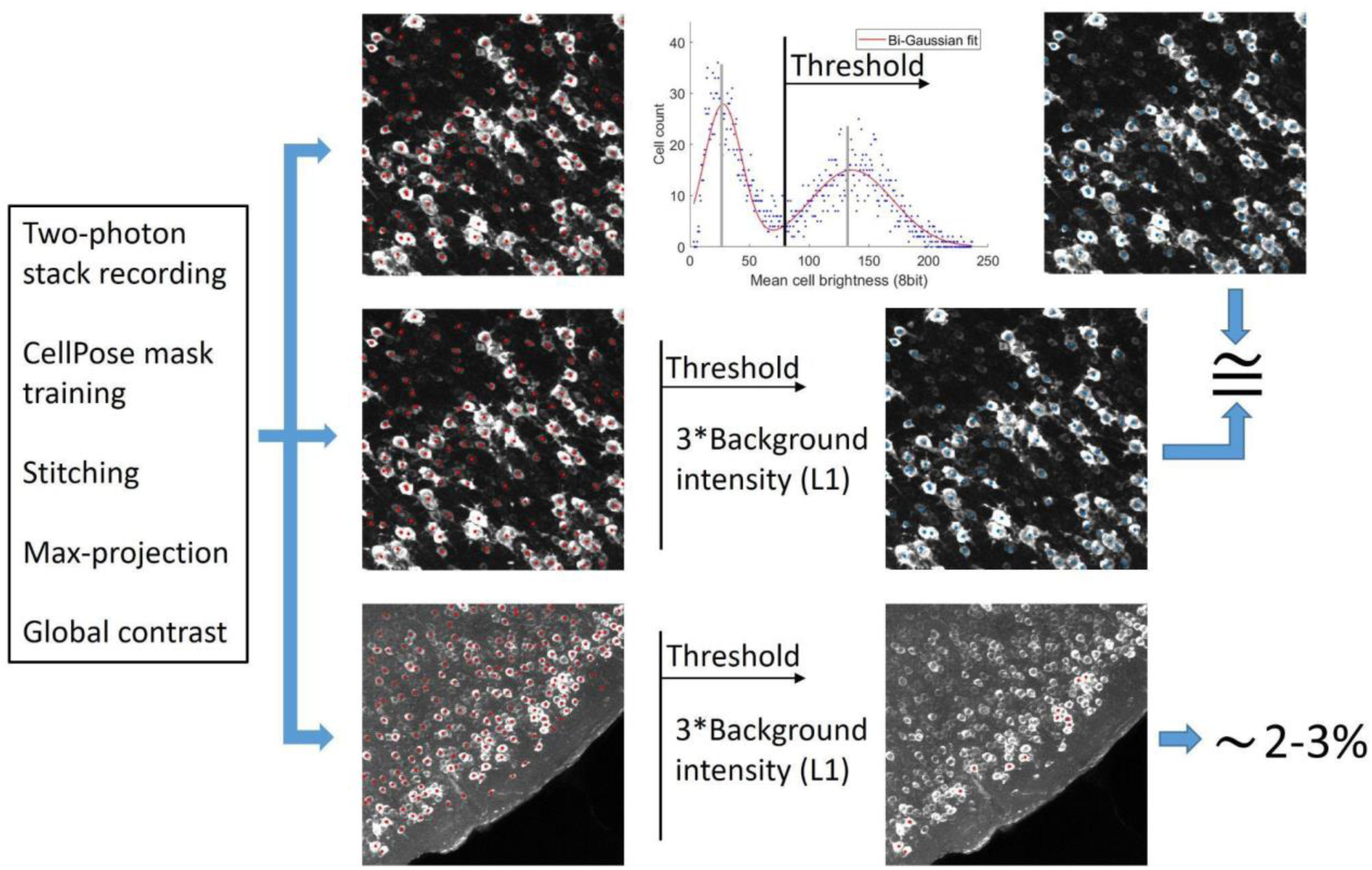
Background-insensitive classification algorithm of pS6-positivity in IHC datasets.

**Supplementary Fig. 3.**
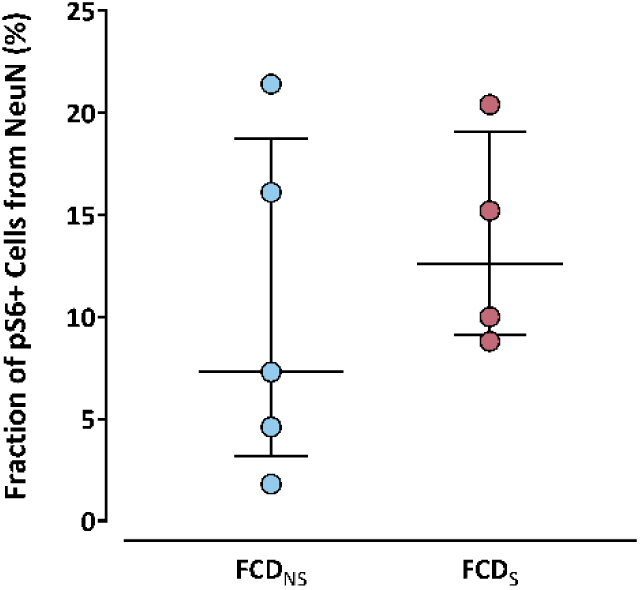
Fraction of mTOR-hyperactivated neurons in FCD in seizing and non-seizing animals. Points denote individual mice, middle lines = medians, whiskers = IQRs.

**Supplementary Fig. 4.**
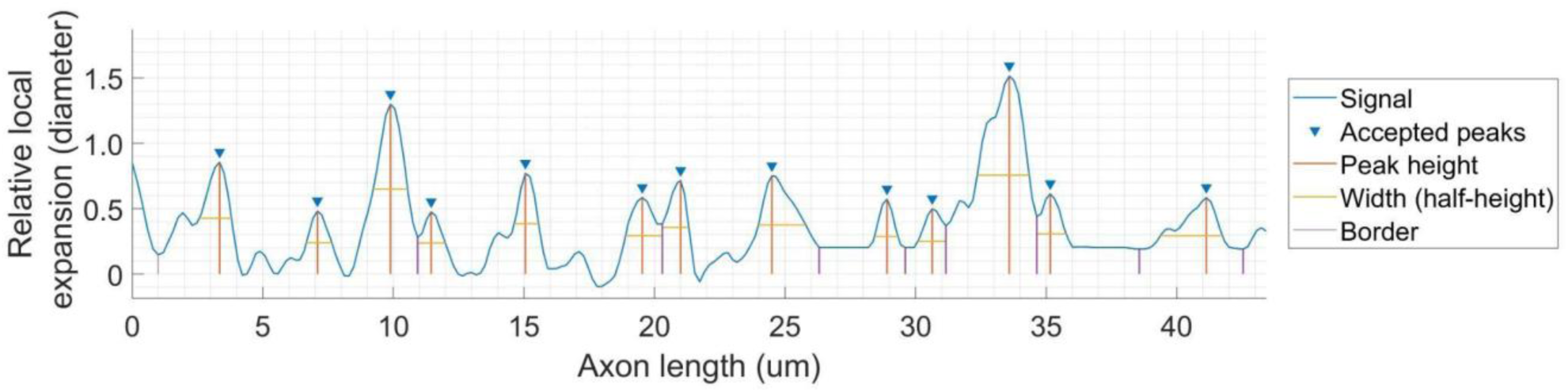
Axonal boutons detection parameters. Individual parameters defined in Methods.

**Supplementary Fig. 5.**
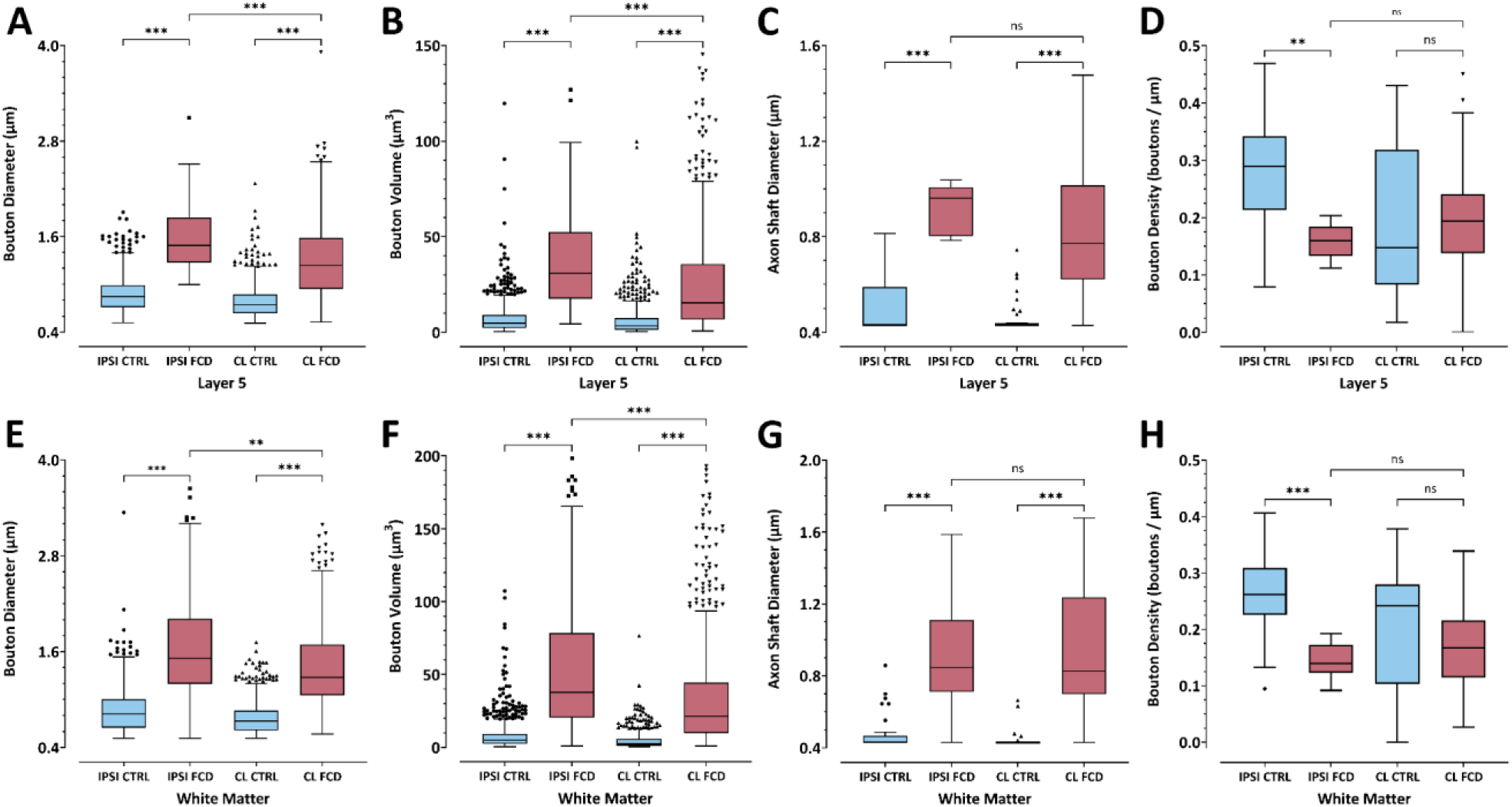
Parameters of ipsilateral and contralateral axonal boutons in layer 5 and white matter. (**A, E**) Tukey boxplots of axonal bouton diameters. (**B, F**) Tukey boxplots of axonal bouton volumes. (**C, G**) Tukey boxplots of axon shaft diameters. (**D, H**) Tukey boxplots of axonal bouton density. (**A–H**) Measurements taken from either layer 5 or the white matter in either control animals (CTRL) or animals with lesions (FCD), both on the ipsilateral (IPSI) and homotopic contralateral (CL) areas.

**Supplementary Tab. 1.** Complete values to Supplementary Fig. 5. Legend: **IC** = ipsilateral control; **IF** = ipsilateral FCD; **CC** = contralateral control; **CF** = contralateral FCD; **IQR** = interquartile interval; **N** = number of boutons/shafts, mult. **p** = p values from multiple comparisons tests.

| Sup. 5<br>Subplot | Median |  |  |  | IQR |  |  |  | N |  |  |  | **P** | **mult. p**<br>(ICvIF, CCvCF,<br>IFvCF) |
| --- | --- | --- | --- | --- | --- | --- | --- | --- | --- | --- | --- | --- | --- | --- |
|  | IC | IF | CC | CF | IC | IF | CC | CF | IC | IF | CC | CF |  |  |
| A (μm) | 0.85 | 1.5 | 0.74 | 1.2 | 0.71–0.99 | 1.3–1.8 | 0.64–0.87 | 0.94–1.6 | 1048 | 115 | 666 | 684 | <0.001 | <0.001 ALL |
| B (μm<sup>3</sup>) | 4.7 | 31 | 3.4 | 16 | 2.3–9.2 | 17–52 | 1.4–7.5 | 6.7–36 | 1048 | 115 | 666 | 684 | <0.001 | <0.001 ALL |
| **C ($\mu\text{m}$)** | 0.43 | 0.96 | 0.43 | 0.77 | 0.43–0.59 | 0.80–1.0 | 0.43–0.44 | 0.62–1.0 | 46 | 9 | 36 | 54 | <0.001 | <0.001,<br><0.001, 0.61 |
| **D ($\mu\text{m}^{-1}$)** | 0.29 | 0.16 | 0.15 | 0.19 | 0.21–0.34 | 0.13–0.18 | 0.082–0.32 | 0.14–0.24 | 46 | 9 | 36 | 54 | <0.001 | 0.002, >0.99,<br>0.52 |
| **E ($\mu\text{m}$)** | 0.82 | 1.5 | 0.73 | 1.28 | 0.65–1.0 | 1.2–2.0 | 0.62–0.86 | 1.1–1.7 | 966 | 237 | 716 | 632 | <0.001 | <0.001,<br><0.001, 0.004 |
| **F ($\mu\text{m}^3$)** | 4.9 | 38 | 2.5 | 21 | 2.3–9.2 | 20–78 | 1.2–5.9 | 9.8–44 | 966 | 237 | 716 | 632 | <0.001 | <0.001 ALL |
| **G ($\mu\text{m}$)** | 0.43 | 0.84 | 0.43 | 0.83 | 0.43–0.47 | 0.71–1.1 | 0.43–0.43 | 0.70–1.2 | 46 | 19 | 38 | 45 | <0.001 | <0.001,<br><0.001, >0.99 |
| **H ($\mu\text{m}^{-1}$)** | 0.26 | 0.14 | 0.24 | 0.17 | 0.23–0.31 | 0.12–0.17 | 0.10–0.28 | 0.11–0.22 | 46 | 19 | 38 | 45 | <0.001 | <0.001, 0.09,<br>0.90 |

**Supplementary Fig. 6.**
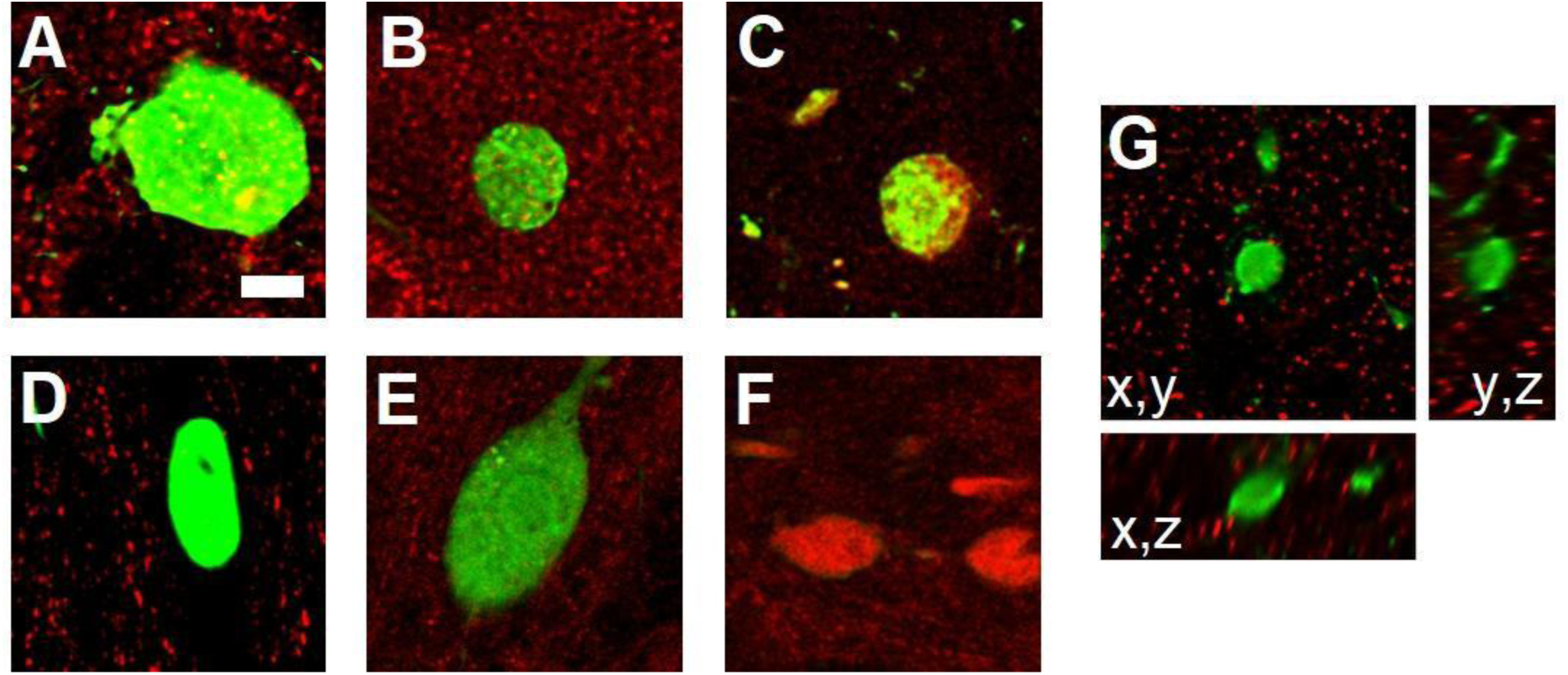
Examples of axonal varicosities in the contralateral cortex and the corpus callosum stained for synapse-related proteins. Giant axonal boutons of dysmorphic neurons were visualized in the contralateral (CL) cortex (**A**–**C**, **G**) and in the corpus callosum (CC) (**D**–**F**). (**A**) vGlut1 puncta (red) colocalized with axonal boutons (green) in the CL cortex, but not in the CC (E). (**B**) Synapsin puncta (red) were present within boutons (green) in the CL cortex, but absent from CC boutons (**E**). (**C**) Synaptic terminals of dysmorphic neurons expressing tdTomato and the synaptophysin–EGFP fusion protein. Clusters of synaptophysin–EGFP (green) were evident in boutons in the CL cortex, but not in axonal varicosities in the CC (**F**). (**G**) An axonal bouton of an EGFP-expressing dysmorphic neuron labeled against PSD95 (red). Scale bars: 5 μm.

